# Characterization of EDS1-independent plant defense responses against bacterial pathogens using Duckweed/*Pseudomonas* pathosystems

**DOI:** 10.1101/2022.03.31.486129

**Authors:** E.L Baggs, M.B Tiersma, B.W Abramson, T.P Michael, K.V Krasileva

**Affiliations:** Department of Plant and Microbial Biology, University of California Berkeley, USA; Plant Molecular and Cellular Biology Laboratory, The Salk Institute for Biological Studies, La Jolla, USA

## Abstract

ENHANCED DISEASE SUSCEPTIBILITY 1 (EDS1) mediates the induction of defense responses against pathogens in most land plants. However, it has recently been shown that a few species have lost EDS1. It is unknown how defense against disease unfolds and evolves in the absence of EDS1. Here we utilize duckweeds; a collection of aquatic species that lack EDS1, to investigate this question. We successfully established duckweed-*Pseudomonas* pathosystems and were able to characterize pathogen-induced responses in an immune system that lacks the EDS1 signaling pathway. We show that the copy number of infection-associated genes and the infection-induced transcriptional responses of duckweeds differ from that of other model species. Moreover, we show that the conservation of canonical Microbe Triggered Immunity and Effector Triggered Immunity pathways varies between duckweed species. This work shows that pathogen defense has evolved along different trajectories and uncovers alternative genomic and transcriptional reprogramming. Specifically, the miAMP1 domain containing proteins, which are absent in Arabidopsis, show pathogen responsive upregulation in duckweeds. Despite such divergence between Arabidopsis and duckweed species, we find evidence for the conservation of upregulation of certain genes and the role of hormones in response to disease. Our work highlights the importance of expanding the pool of model species to study defense responses that have evolved in the plant kingdom, including those independent of EDS1.

## Introduction

Receptors and signaling components of the plant immune signal network likely evolved before the divergence of flowering plants 181 million years ago (MYA) (Kumar et al. 2017). The two primary layers of plant innate immunity, Microbe Triggered Immunity (MTI) and Effector Triggered Immunity (ETI) (Alhoraibi et al. 2019), form an intricate network of cross-amplifying pathways (Ngou et al. 2021; Yuan, Jiang, et al. 2021; Tian et al. 2020). The shared evolutionary history of MTI and ETI suggests that they intricately co-evolved and are foundational for immunity in flowering plants. ENHANCED DISEASE SUSCEPTIBILITY 1 (EDS1) is a central hub in MTI and ETI signal transduction and amplification. Mutations abolishing activity of EDS1 compromise ETI and MTI (Pruitt et al. 2021; Cui et al. 2017; Wagner et al. 2013). Surprisingly, genomic studies have revealed a number of land plants that thrive in their natural environments without EDS1, including the Lemnaceae (E. L. Baggs et al. 2020). It is unknown how the response to bacterial pathogens would proceed or evolve in the absence of a functional EDS1.

Lemnaceae diverged from other monocots 128 MYA (Kumar et al. 2017). The common ancestor of the Lemnaceae was present in a freshwater aquatic environment and had a reduced body plan of a frond (a fused stem and leaf) and root (Acosta et al. 2021). Genera within the Lemnaceae include *Spirodela, Landoltia, Lemna, Wolffia* and *Wolffiella*, all of which primarily reproduce through asexual budding (Bog, Appenroth, and Sree 2019). The small genome and body plan is conducive to the Lemnaceae’s rapid lifecycle with a doubling time of as little as 34 hours (Michael et al. 2020). Lemnaceae are very easy to grow in the laboratory which has stimulated research into their use as biofuels (Van Hoeck et al. 2015; Y.-L. Xu et al. 2018; Su et al. 2014) and enabled the growth of genomic resources for the species, including multiple genomes and transcriptomic datasets (Michael et al. 2020; P. T. N. Hoang et al. 2020; W. Wang et al. 2014; Michael et al. 2017; Wenguo Wang et al. 2016; Abramson et al. 2021). *Spirodela polyrhiza* has a relatively small genome of 158 Mb with less than 20,000 protein coding genes and has lost members of expansin and cellulose biosynthesis (W. Wang et al. 2014; Michael et al. 2017)*. Wolffia australiana* has ~15,000 protein coding genes having lost light signaling pathways and root development pathways, which is not surprising given the absence of roots in *Wolffia* (Michael et al. 2020). In addition to developmental pathways lost in other Lemnaceae, *S. polyrhiza* was previously shown to have lost *EDS1* (E. L. Baggs et al. 2020), however the extent of immune pathway loss in other Lemnaceae remained unclear.

Given the absence of the EDS1 pathway in *S. polyrhiza* (E. L. Baggs et al. 2020), we hypothesized that duckweeds could be used as a model for understanding EDS1-independent innate immunity. Lemnaceae are globally distributed and only absent from the Arctic and Antarctica (Landolt 1992; Crawford et al. 2006); several Lemnaceae species are even noted in the invasive species compendium (“Invasive Species Compendium” n.d.) (Abramson et al. 2021). The wide distribution of duckweeds means their range overlaps that of a number of model plant pathogens. These include *Pseudomonas syringae pv. tomato* DC3000, *Pseudomonas syringae pv. syringae, Xanthomonas perforans, Xanthomonas euvesicatoria* (Potnis et al. 2015; Cai et al. 2011; Gutiérrez-Barranquero, Cazorla, and de Vicente 2019), and many others for which genomic and genetic resources are available. These pathogens often have devastating effects on crop plants yield and value (Martins et al. 2018).

MTI immune signaling is triggered by the recognition of conserved molecular patterns through Pattern Recognition Receptors (PRR) and their respective co-receptors. There are two main types of PRRs: Receptor-Like Kinases (RLKs) and Receptor-Like Proteins (RLPs), that lack an intracellular kinase domain (Offor, Dubery, and Piater 2020; Yuan, Ngou, et al. 2021; Lolle, Stevens, and Coaker 2020). Substantial differences exist between RLP and RLK signaling pathways: immune signaling through the RLP and Suppressor of BIR1 (SOBIR1) co-receptor pathway genetically requires EDS1, Phytoalexin Deficient 4 (PAD4) and Resistance to Powdery Mildew 8 NLRs (RNLs) and results in higher levels of ethylene and phytoalexins production, as well as Pathogenesis Related 1 gene expression (Tian et al. 2021; Pruitt et al. 2021). Pathogenesis Related (PR) genes are characterized by their rapid upregulation after pathogen infection, they include a number of antimicrobial genes with different modes of action (glucanase, thaumatin, chitinase, thionin and defensin) (Ali et al. 2018). There are many antimicrobial peptides that are not classed as PR genes such as proteins typified by the presence of the MiAMP1 domain (McManus et al. 1999; Marcus et al. 1997; Stephens et al. 2005), but the diversity and modes of action of these antimicrobials remains poorly understood.

RLP and RLK activation primes the cell to initiate a stronger immune response upon Nucleotide-binding Leucine-rich repeat Receptors (NLRs) perception of intracellular changes caused by pathogen-derived effectors (Tian et al. 2020). Conserved domain architecture and signaling specificities of NB-ARC domain containing proteins allow their sub-categorisation into RNLs, coiled-coil NLRs (CNLs), Toll/interleukin-1 receptor NLRs (TNLs) and TIR-NBARC-like-β-propeller (TNP) (Nandety et al. 2013; E. Baggs, Dagdas, and Krasileva 2017; Shao et al. 2019, 2016; Johanndrees et al. 2021). Disease resistance mediated by TNLs and some CNLs is genetically dependent on RNLs and the lipase-like proteins EDS1, PAD4 and SAG101. The recognition of pathogen presence by an NLR typically leads to qualitative resistance where the plant shows a discrete resistant phenotype.

In the absence of NLR triggered ETI plants are not necessarily susceptible to a pathogen instead quantitative resistance may be observed where extent of resistance is more variable and in some cases underpinned by hundreds of loci (Corwin and Kliebenstein 2017). Mechanisms implicated in quantitative resistance include Defensins, pathogenesis-related proteins, secondary metabolite enzymes and pathogen-induced phytohormones accumulation (Corwin and Kliebenstein 2017).

Qualitative disease resistance results in mendelian patterns of inheritance of discrete resistant and susceptible individuals when investigating a given pathogen genotype. Typically the mechanism of qualitative resistance involves MTI and ETI. In contrast, quantitative resistance underpins a continuous distribution in the response among related individuals to a pathogen genotype, in this case resistance is governed by a number of small effect loci and doesn’t necessarily require MTI/ETI pathways. The dynamics of plant-pathogen interactions are complex and can be manipulated by both the plant and pathogen to their own benefit

In this study, we investigated the phenotypic, genomic and transcriptomic characteristics of duckweeds in response to *Pseudomonas* plant pathogens. We found a stepwise reduction of conserved ETI pathways within the Lemnaceae and loss of MTI immune pathway components in *W. australiana*. On the other hand, we observed a shared expansion of the MiAMP1 protein family across Lemnaceae. Additionally, we observed species specific responses to pathogen treatments among duckweed species. We investigated the transcriptional response to pathogens of duckweed species, a system which is adapted to the absence of EDS1-mediated immune signaling cascades. Our study highlights duckweeds as a rapid growth, high throughput, minimalist MTI-ETI model organism. As such, it could be utilized to expedite our understanding of EDS1-independent MTI-ETI mechanisms of immunity as well as to dissect mechanisms of quantitative resistance.

## Results

### Lemnaceae retained most conventional MTI signaling components and expanded the MiAMP protein family

To understand how the repertoire of immune genes has diverged within the Lemnaceae, we utilized publicly available genomes of *S. polyrhiza* and *W. australiana*. Additionally, we assembled and annotated the genome of *Landoltia punctata* clone 5635, which allowed us to catalog the presence and absence variation of immunity genes across the 3 duckweed genera (Fig 1a, S.table 1). We found that *EDS1, PAD4*, and *RNLs* were absent across the Lemnaceae, consistent with previous findings (Lapin et al. 2019; E. L. Baggs et al. 2020; Michael et al. 2020). To estimate the timing of the EDS1 pathway loss, we leveraged giant Taro (*Colocasia esculenta*), which is the closest related species to duckweeds with an available genome (Yin et al. 2021). Taro has retained the *EDS1* pathway genes, suggesting that the loss of these genes in Lemnaceae occurred after divergence of Lemnaceae and Taro 104-117 MYA (Kumar et al. 2017).

**Figure 1.**
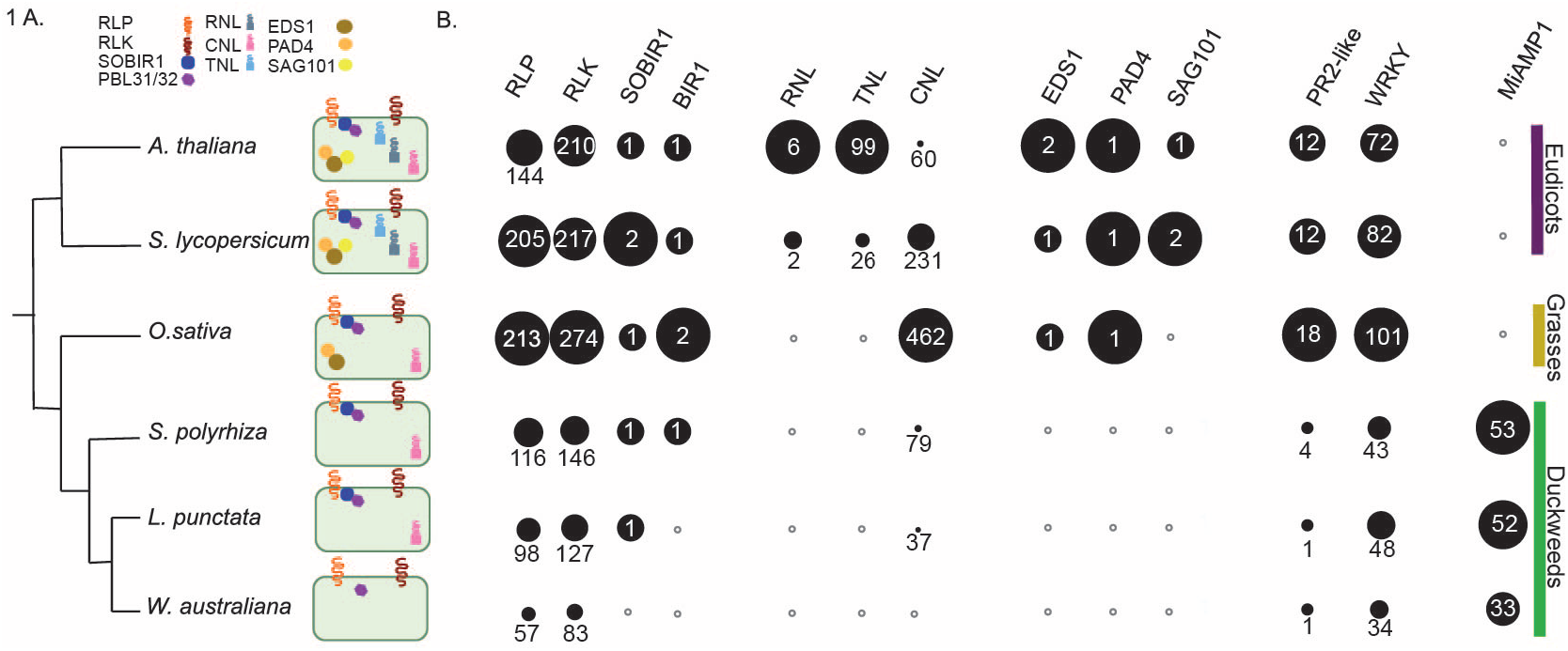
Copy number variation of conserved angiosperm immune signaling components. A. Phylogenetic relationship of duckweeds to other representative angiosperms and depiction of presence of signaling components. If components were not identified by Orthofinder, reciprocal BLASTP, and TBLASTN, the representative symbol was not displayed in the cell. B. The size of circles within each column are proportional to the highest copy number of that gene family in a given species, the copy number is denoted in white text. White circles with a gray outline indicate no members of that gene family were identified.

To understand the extent to which immune signaling pathways diverged across Lemnaceae, we surveyed the presence of 24 additional protein families with a known role in plant disease defense and which arose prior to monocot and dicot divergence. All of the 24 additional protein families we selected were present in *S. polyrhiza* (S.table 1,2, Sfig 1-3). Though gene families associated with immunity were often retained in the Lemnaceae, we noticed a reduction in copy number similar to the reduced copy number of NLRs (Fig 1b, Sfig 1, S.table 1). *W. australiana* has only 3 NB-ARC domain-containing proteins (Michael et al. 2020): two of which consists of only the NB-ARC domain and the other is a TNP and its conservation is consistent with the finding that TNPs are EDS1 independent (Johanndrees et al. 2021). Despite this, *W. australiana* has retained many other immune components that we investigated (Fig 1b). While we identified no orthologs of *Pathogenesis related 4 (PR4), SOBIR1* and *BAK1 interacting receptor 1 (BIR1*) in *W. australiana*, they were present in *L. punctata* and *S. polyrhiza* (Fig 1b). Our observations indicate that immune pathways have undergone varying degrees of gene loss across the duckweed genera.

Despite the general trend of reductionism of immune signaling components in the Lemnaceae, we identified the striking expansion of the MiAMP1 type antimicrobial proteins. It has previously been shown that antimicrobial proteins in general expanded in *S. polyrhiza* 7498 (An et al. 2019). Phylogenies of proteins from the MiAMP1 family show lineage specific expansions and rapid birth and death among the Lemnaceae (Sfig 4) suggesting that selection pressures may be favoring the expansion and diversification of MiAMP1 proteins in the duckweed lineage.

### Duckweed species show variable symptoms upon *Pseudomonas* and *Xanthomonas* challenge

Since Lemnaceae species lack the EDS1 pathway, it was unclear how they would respond to bacterial pathogen infection. We therefore challenged *S. polyrhiza, L. punctata*, and *W. australiana* to a panel of common *Pseudomonas* and *Xanthomonas* plant pathogens with a standard high bacterial inoculum (1×10^8^CFU/ml) (Fig 2a) to assess their susceptibility and disease phenotypes. We observed variability among replicates in the susceptibility and resistant phenotypes (Fig 2b, S.table 3, Sdataset 1) which is consistent with quantitative resistance. We identified a few virulent pathogens that produced similar symptoms across all hosts (*Xanthomonas gardneri, Pseudomonas syringae pv. tabaci*) while others caused distinct symptoms on a given host species. The most common disease symptoms we observed were chlorosis and reduced growth rate. Surprisingly, despite having fewer NLRs and lacking SOBIR1 and BIR1, the growth of *W. australiana* upon pathogen infection was often less affected than that of other duckweed species.

**Figure 2.**
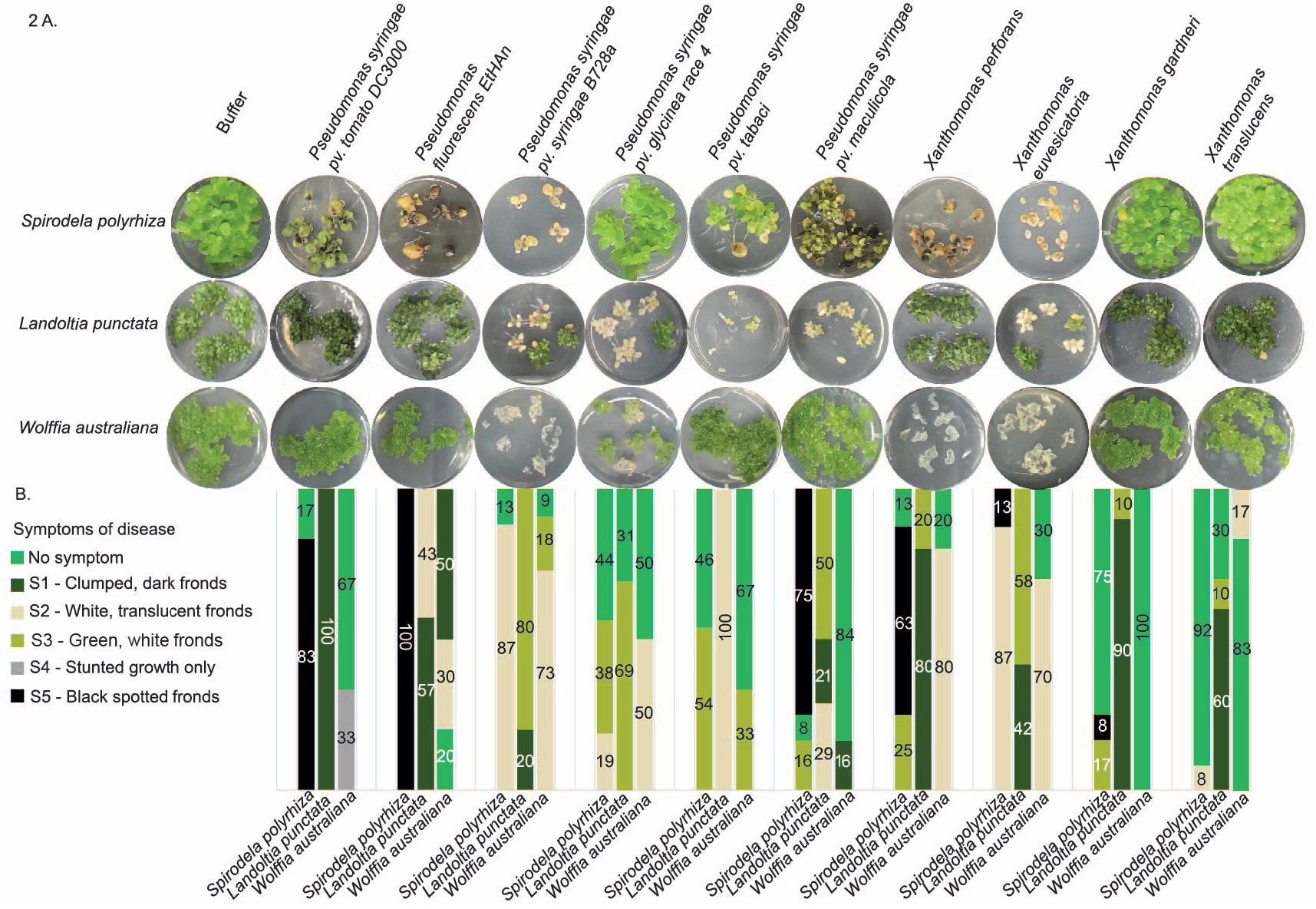
Phenotypic response of duckweed species to bacterial pathogen treatments. A. One replicate is shown per treatment (full experiment has been independently replicated 3 times, see Sdataset 1, S.table 3). Each well displays the most prevalent visual symptomof infection three weeks after pathogen treatment of five frond clusters. B. Bar graph showing the percentages of wells displaying each symptom among all replicates (S.table 3). Colors of oars correspond to color legend of disease symptoms.

For further experiments, we focused on *P. syringae pv. tomato* DC3000 (*Pst* DC3000) and *P. syringae pv. syringae* B728a (*Pss* B728a) as there is detailed knowledge about their virulence factors, large resources of mutants, and the severity of disease symptoms that they caused varied across duckweeds.

### *P. syringae* colonizes the substomatal cavity in *S. polyrhiza* but infection is slowed in T3SS mutants

To further characterize the pathology of *Pseudomonas* on duckweed, we used microscopy and bacterial genetics, taking advantage of the type III secretion deficient mutant *Pst* DC3000 hrcC^-^.

Microscopy of flood inoculated *S. polyrhiza* fronds with standard high bacterial inoculum at 5 days post inoculation (5 dpi) showed the accumulation of GFP expressing *Pst* DC3000 concentrated at the node where the root and frond join and budding pockets were daughter fronds emerge (Sfig 5). By 7 dpi *Pst* DC3000 populations were observed at the stomata and within the substomatal cavity and mesophyll (Fig 3a, Sdataset 2). We also observed *Pst* DC3000 populations on root tissue at 7 dpi (Fig 3b). Together, our observations suggest that the visual symptoms of *Pseudomonas* on duckweed are a result of active bacterial infection. Therefore, the duckweed-*Pseudomonas* pathosystem constitutes a valuable model for understanding disease progression and EDS1-independent immune responses.

**Figure 3.**
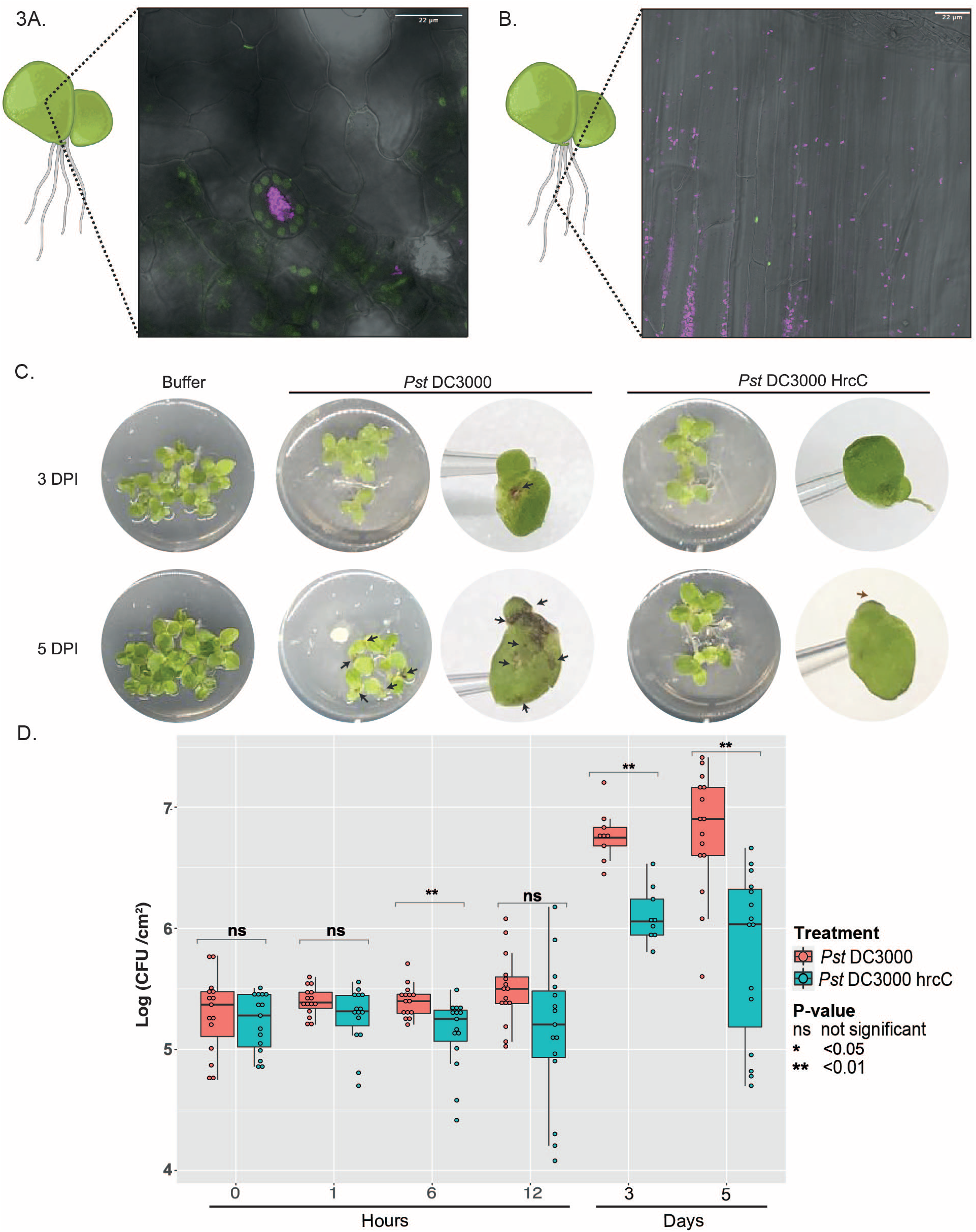
Pst DC3000 infection of Spirodela polyrhiza. Confocal 100x microscopy of *S. polyrhiza* inoculated with Pst DC3000. False colors: pink - *Pst* DC3000, green - plastids, gray - transmitted light. A. Frond surface with stomata in the center of frame B. Root tissue. C. Images of duckweed fronds and symptoms during *Pst* DC3000 and *Pst* DC3000 hrcC^-^. The *Pst* DC3000 hrcC^-^ mutant lacks the type III secretion system used for deployment of effectors to the plant. Individual fronds were photographed on a 10ul tip to indicate size of fronds. D. Box plot showing the number of colony forming units of *Pst* DC3000 and *Pst* DC3000 hrcC^-^ at different time points on *S. polyrhiza* fronds. Dots indicate individual replicates.

To advance our understanding of this pathosystem, we infiltrated *S. polyrhiza* plants with *Pst* DC3000 and *Pst* DC3000 hrcC^-^ with a low bacterial inoculum (1×10^5^ CFU/ml), and monitored bacterial population growth overtime (Fig 3c/d). The first black lesions were macroscopically visible at 3 days post inoculation with *Pst* DC3000 and by day 5 we observed an increase in the size, number of lesions and proportion of affected fronds (Fig 3c). Consistent with the dense populations at the budding pocket and node observed by microscopy, the macroscopic black lesions observed were initially localized at the budding pocket and node. We observed no black lesions upon inoculation with *Pst* DC3000 hrcC^-^ at 3 or 5 dpi, although the frond edges appeared shriveled at 5 dpi. In contrast, *Pst* DC3000 or *Pss* B728a infiltrated *L. punctata* even 5 days post inoculation showed no macroscopically visible lesions (Fig 4a/b). 3 weeks post inoculation, 5/8 replicates of *L. punctata* inoculated with *Pss* B728a had turned white and produced fewer daughter fronds than in other treatments. However, even at 3 weeks post inoculation we observed no severe symptoms for *Pst* DC3000, although the fronds showed slight dark discoloration and clumping. *Pst* DC3000 hrcC^-^ did not cause any symptoms on *L. punctata* throughout the 3 weeks post inoculation (Sfig 6).

**Figure 4.**
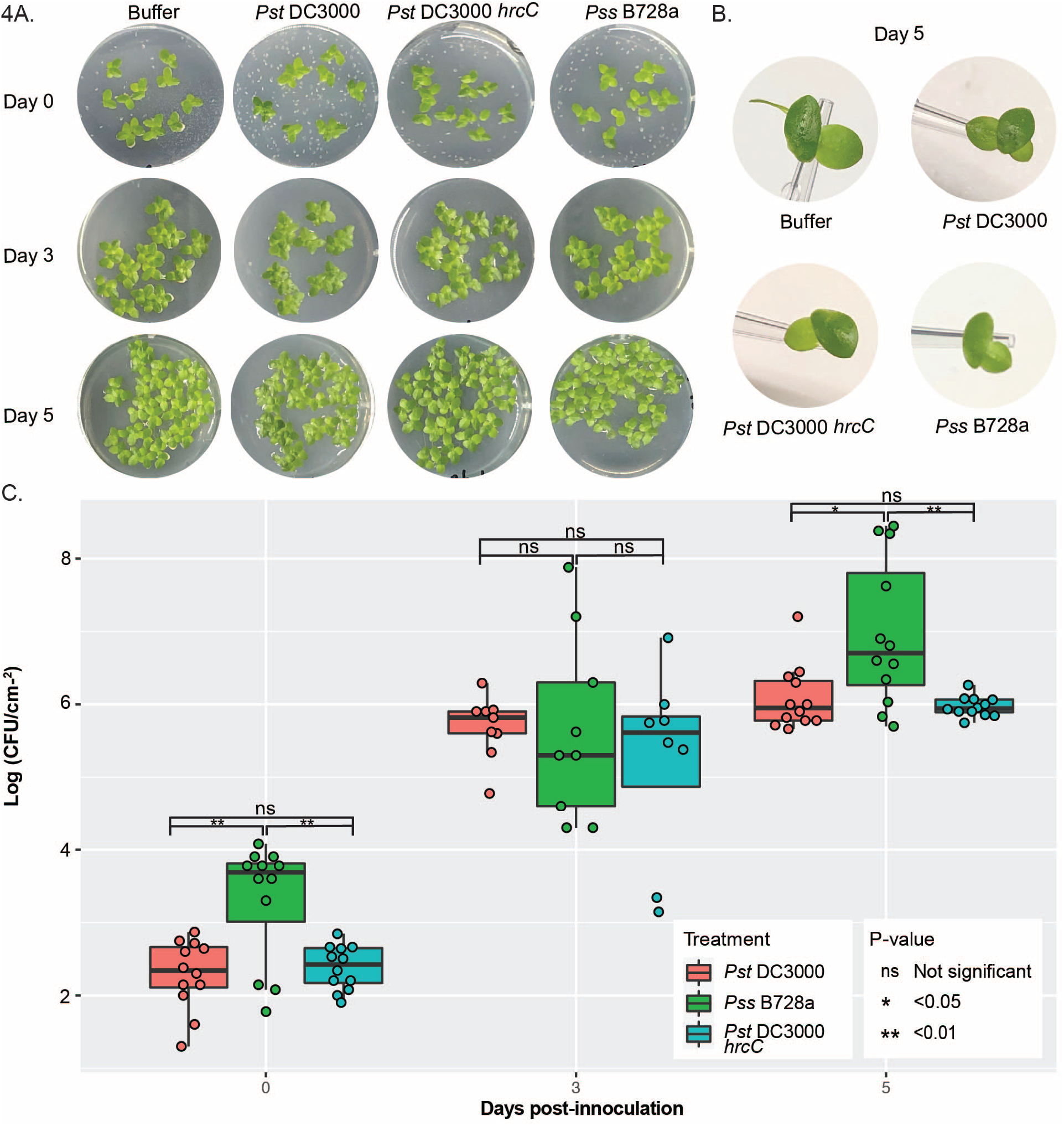
*Pst* DC3000 infection of *Landoltia punctata*. A. Images of *L. punctata* fronds and symptoms during *Pst* DC3000, *Pst* DC3000 hrcC^-^ and *Pss* B728a.B. Individual fronds were photographed on a 10ul tip to indicate size of fronds. C. Box plot showing the number of colony forming units of *Pst* DC3000, *Pst* DC3000 hrcC^-^ and *Pss* B728a at different time points on *L. punctata* fronds. Dots indicate individual replicates. Mean-T statistic with holm adjustment used to asses statistical difference between treatments.

In addition to causing visible disease symptoms, *Pst* DC3000 and *Pst* DC3000 hrcC^-^ were able to proliferate within *S. polyrhiza* (Fig 3d). *Pst* DC3000 multiplied to similar levels as in previously described compatible interactions with wild type *A. thaliana* (Ishiga et al. 2011; Velásquez et al. 2017; Katagiri, Thilmony, and He 2002). However, we observed significantly fewer colony forming units of *Pst* DC3000 hrcC^-^ compared to *Pst* DC3000. This suggests that the virulence of *Pst* DC3000 on *S. polyrhiza* relies on the presence of effectors and the type III secretion system. In contrast, consistent with the lack of disease symptoms in *L. punctata* neither *Pst* DC3000 nor *Pst* DC3000 hrcC^-^ showed significantly different CFU counts at 5 dpi. Instead, both bacterial numbers remained at a level similar to those of *Pst* DC3000 hrcC^-^ in *S. polyrhiza* (Fig 4c). Interestingly, although we saw no visible lesions on *L. punctata* with *Pss* B728a 5 days after inoculation, the CFU counts were significantly higher than those of *Pst* DC3000 after 5 days but were similar to those of *Pst* DC3000 in *S. polyrhiza*. Our results show that *Pseudomonas* pathogens *Pst* DC3000 and *Pss* B728a are able to proliferate in duckweed host and bacterial multiplication is affected by unknown host factors and for *S. polyrhiza / P*st DC3000 interaction the type III secretion system.

### Lemnaceae spp. and pathogen treatment affect outcome of hormone treatment

In response to biotrophy, plants upregulate the phytohormones salicylic acid (SA) (Cao et al. 1994; Clarke et al. 1998) and N-hydroxy pipecolic (NHP) through pathways dependent and independent of EDS1 which amplifies defense response signaling cascades (Chen et al. 2018). To suppress phytohormone responses, *Pst* DC3000 produces the toxin coronatine (Moore et al. 1989; Mittal and Davis 1995), a structural mimic of jasmonic acid which counteracts SA upregulation (Fonseca et al. 2009; Wasternack and Xie 2010). Since *Pst* DC3000 was virulent on *S. polyrhiza*, we investigated the role of phytohormones and toxins in the absence of EDS1 through our duckweed pathosystem. The *Pst* DC3000 *cma*^-^ mutant is deficient in coronatine biosynthesis and causes mild symptoms on *S. polyrhiza*, marked by the absence of black lesions (Fig 5a, Sfig 7,8). Addition of coronatine alone (0.3 μM) was sufficient to disfigure fronds, reduce growth, and sometimes induced the formation of turion resting bodies. The effect on growth and turion formation was stronger at higher concentrations of coronatine (3 μM). However, addition of coronatine on its own was not sufficient to recover symptoms of black lesions and white bleaching of fronds comparable to *Pst* DC3000 infection. The treatment of fronds with coronatine at the time of *Pst* DC3000 *cma^-^* infection restored the black lesions to some extent, but the strain still did not induce white bleaching. Our results show that coronatine has a conserved role in promoting pathogen virulence in duckweeds despite the absence of the SA promoting EDS1 pathway.

**Figure 5.**
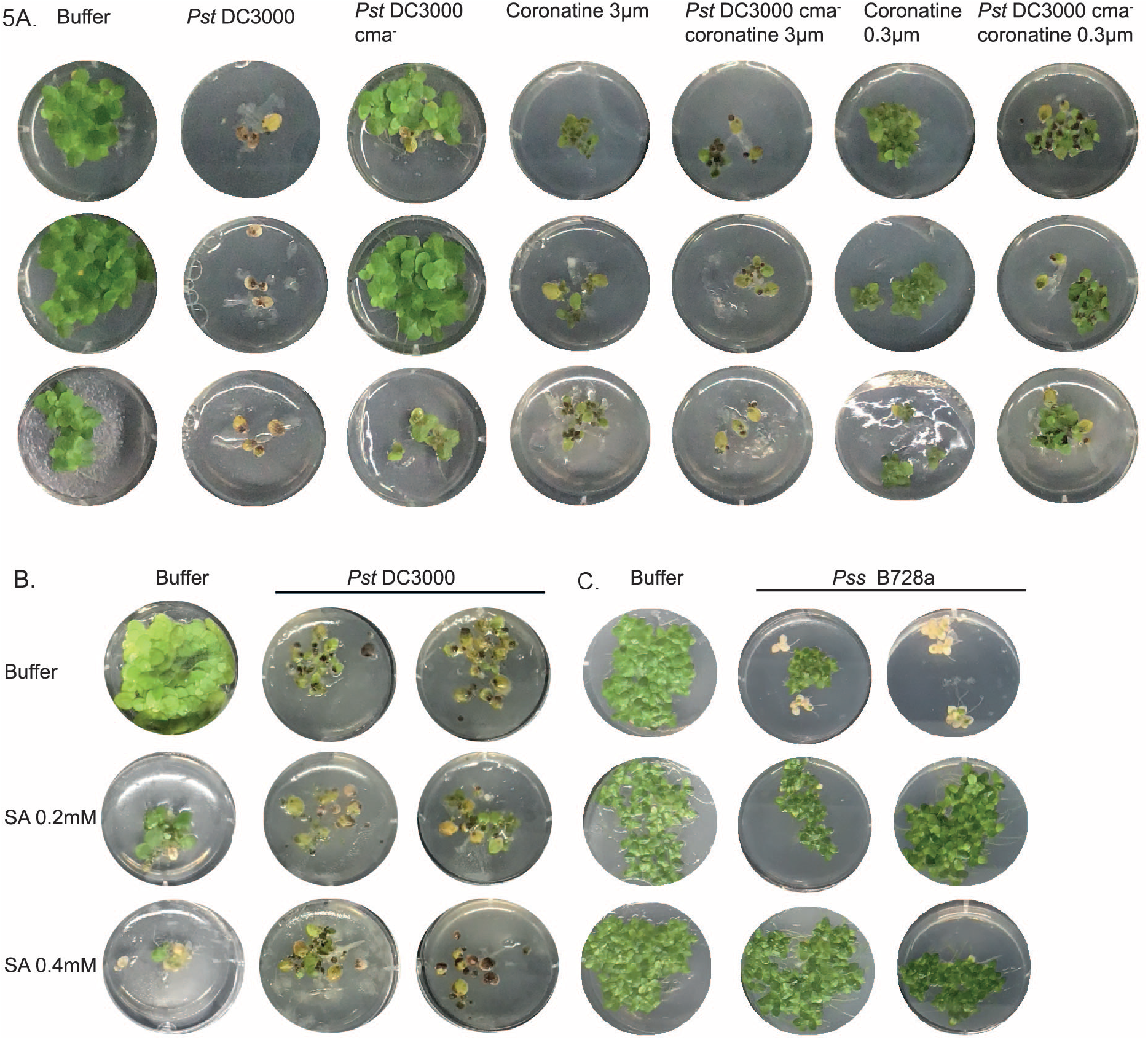
The effects of hormone treatments on *Pst* DC3000 and *Pss* B728a disease symptoms. A. Images of disease progression 3 weeks after the treatment of 3*S. polyrhiza* fronds with different concentrations and combinations of coronatine and *Pst* DC3000 mutants (full experiment has been independently replicated 3 times, see Sfig 7 and Sfig 8). B. Images of disease progression 3 weeks after the treatment of 5 *S. polyrhiza* fronds with different concentrations and combinations of salicylic acid and *Pst* DC3000 (full experiment has been independently replicated 4 times, see Sfig 10-12). C. Images of disease progression 3 weeks after the treatment of five *L. punctata* fronds with different concentrations and combinations of salicylic acid and Pss B728a treatment, Pss B728a was used rather than *Pst* DC3000 as it gives clearer visual symptoms in *L. punctata*, (full experiment has been independently replicated 3 times, see Sfig 14 and Sfig 15).

Given that a core function of the EDS1 pathway is to reinforce SA signaling by inducing expression of several SA responsive genes (Jirage et al. 1999; Bonardi et al. 2011; Cui et al. 2017; Roberts et al. 2013); we were interested in how exogenous SA treatment of Lemnaceae would affect *Pst* symptoms. It has been observed in other model plants that amplification of SA can be protective against symptomatic *Pst* DC3000 infection (Tsai et al. 2011). Additionally, SA is perceived and regulated by members of the NPR gene family. Lemnaceae, however, have fewer *NPR* genes than most other species which makes their SA-associated response to phytopathogens of interest. Treatment of Lemnaceae with high levels of SA (2 mM) that are tolerated in *Arabidopsis* were phytotoxic to *S. polyrhiza* (Sfig 9). We therefore carried out experiments using lower levels of SA of 0.4 mM and 0.2 mM. Even at these lower concentrations, we observed dose-dependent frond disfiguration and growth reduction (Fig 5b, Sfig 10-12). As mentioned before, in the absence of SA we observed variation in the virulence of *Pst* DC3000 on duckweed (Fig 2) and this variation translated to the ability of SA to provide pathogen protection. *S. polyrhiza* co-treated with SA and *Pst* DC3000 showed typical *Pst* DC3000-induced symptoms (Fig 5b, Sfig 10-12). However, a small protective effect of SA was observed when *Pst* DC3000 appeared to be less virulent (Sfig 13). We hypothesize that the variation in virulence of *Pst* DC3000 affects the ability of SA to restrict pathogen growth below a critical threshold. In contrast, in the *L. punctata / Pss* B728a pathosystem, there was a clearer protective effect of SA. Priming of *L. punctata* with SA resulted in complete loss of the chlorotic symptoms characteristic of the *Pss* B728a treated fronds (Fig 5c, Sfig 14,15). The effect of SA priming on the outcome of duckweed pathogen interactions appears to be specific to pathogen strain and plant genotype and further complicated by the strong endogenous effect of SA on duckweed physiology.

### *S. polyrhiza and L. punctata* mount transcriptional responses to *Pst* DC3000

Next, we decided to investigate the transcriptional response of duckweeds to *Pseudomonas* infection. Despite the absence of *EDS1*, RNA-seq for both *S. polyrhiza* and *L. punctata* revealed a substantial transcriptional response as early as 30 minutes post infection (Fig 6a, Sfig 16, Sdataset 3, 4). We first examined gene families that are differentially expressed upon pathogen treatment in other plant species: WRKY (Dong, Chen, and Chen 2003), MiAMP1 (Fig 6b) (Adomas and Asiegbu 2006; Adomas et al. 2007), NB-ARC (Richard et al. 2018), and JAZ domain-containing genes (Ishiga et al. 2013) (Sfig 17, S.table 4-11). While we did not observe consistent upregulation Log_2_FC >1 of NB-ARC and WRKY domain-containing genes, MiAMP1 domain-containing genes were consistently upregulated over time in both duckweed species after pathogen treatment.

**Figure 6.**
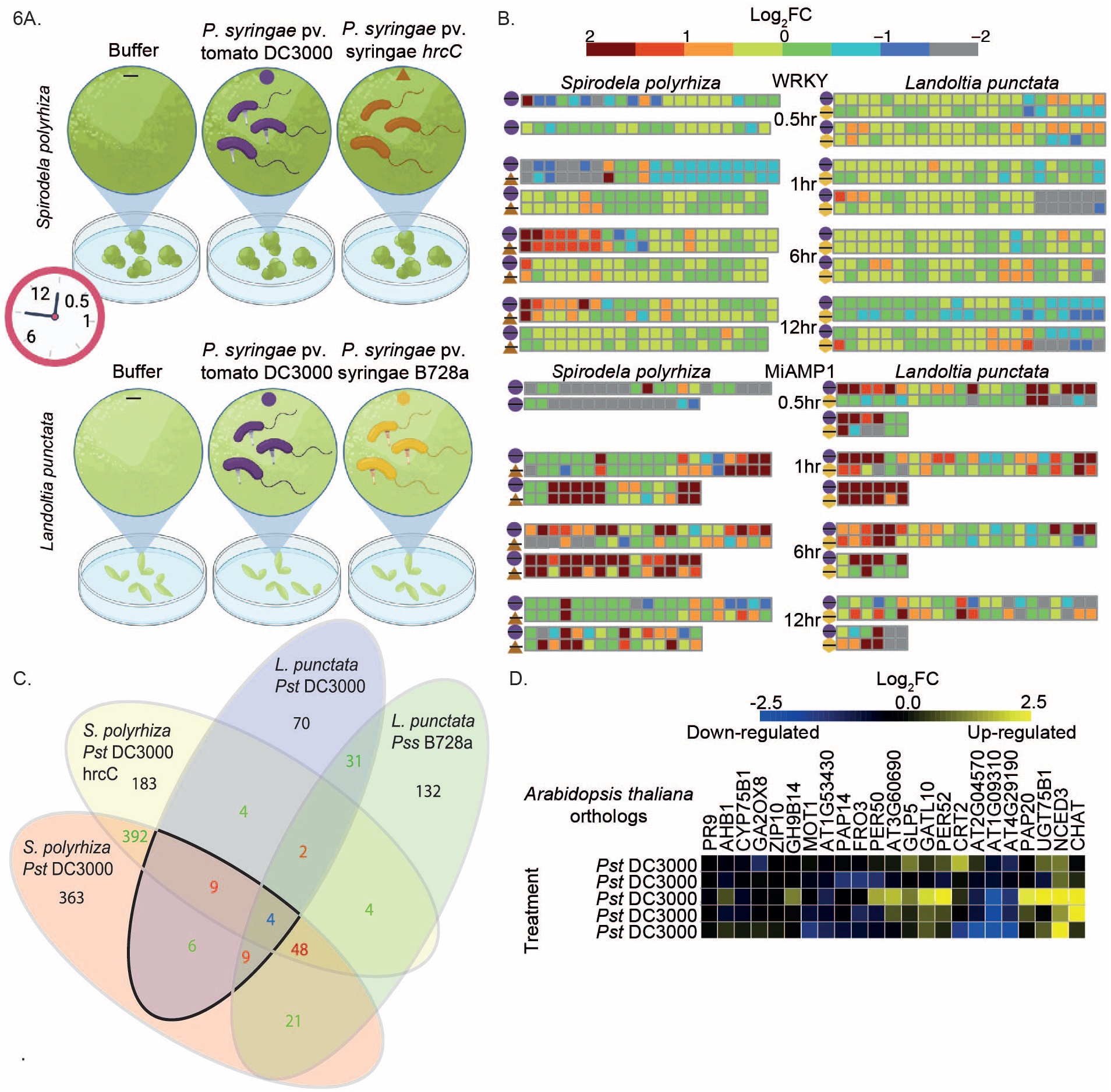
Log_2_ fold change of selected gene families following bacterial pathogen exposure of duckweeds. A. Schematic of the sampling regime for RNAseq study. B. Each square represents a gene with a given domain, the differential expression of the gene is shown by the color of the square. The treatment comparison is indicated by the shapes shown in A. C.Venn diagram of numbers of orthogroups that show differential expression (Log2 FC >1, FDR < 0.05) in multiple combinations of pathogen treatments and or across different species at any time point. D. Microarray differential expression analysis of representatives from *Arabidopsis thaliana* Col-0 of those orthogroups with conserved upregulation upon *Pst* DC3000 in *S. polyrhiza* and *L. punctata*.

We hypothesized that there may be some orthologs of *A. thaliana* bacterial-response genes that are also upregulated in response to bacterial infection in duckweed species. Among the 33 *S. polyrhiza* and 32 *L. punctata* genes considered homologous to *A. thaliana* bacterial-responsive genes (Bjornson et al. 2021; Salguero-Linares et al. 2021; Boudsocq et al. 2010) (S.table 12,13, Sfig 18), we were able to identify 5 *S. polyrhiza* and 5 *L. punctata* gene orthogroups that were significantly upregulated upon pathogen treatment ((Log_2_FC > 1, FDR < 0.05) S.table 14). To investigate whether this conservation extends further we used orthofinder (Emms and Kelly 2019) to create interspecific orthogroups. We then identified orthogroups with members that were significantly differentially expressed in both duckweed species after *Pst* DC3000 treatment (Fig 6c, S.table 15, Sdataset 5). For orthogroups with conserved upregulation in both duckweed species, we investigated if the *A. thaliana* homologs were similarly upregulated upon *Pst* DC3000 infection, using publicly available RNAseq datasets (Fig 6d). Among the *A. thaliana* homologs, 8 genes showed consistent upregulation after *Pst* DC3000 treatment, including a superoxide dismutase *GERMIN-LIKE PROTEIN 5 (GLP5*). In contrast, *PATHOGENESIS RELATED 9 (PR9*) showed conserved upregulation in duckweed but was not upregulated in *A. thaliana* upon *Pst* DC3000 treatment. There appears to be limited overlap in the transcriptional response to *Pst* DC3000 between *A. thaliana* and duckweeds. Interestingly, half of the orthogroups with conserved upregulation in response to bacterial infection were hormone-regulated or biosynthetic genes suggesting a conserved role of hormones in duckweeds and Arabidopsis.

Finally, our analysis allowed the identification of genes whose upregulation was unique to duckweed. Four orthogroups which showed conserved differential expression in both duckweed species do not have homologs in *A. thaliana* (S.table 15). Two of these orthogroups were cytochrome P450 domain-containing genes and a third orthogroup contained enzymes involved in secondary metabolite modifications. Cytochrome P450s are present in *A. thaliana*; however, clades of these proteins appear to have diversified further since the divergence of monocots and dicots (Nelson et al. 2004). MiAMP1 domain-containing genes are not present in *A. thaliana* and although they were upregulated across timepoints in both duckweed species. However, no orthogroup containing these proteins met the criteria for conserved Lemnaceae upregulation. Duckweed species do not show a qualitative but a quantitative resistance phenotype against *Pseudomonas* pathogens and MiAMP and cytochrome P450s proteins could contribute towards this observed quantitative resistance.

## Discussion

Plant immune systems are complex networks of signaling cascades; the crosstalk within the network can mask underlying nodes and their interdependencies. EDS1 is a hub for such signaling and crosstalk. In this study, we investigated the immune response of Lemnaceae species whose immune pathways have evolved for ~110 million years in the absence of *EDS1*. Whilst the loss of *EDS1* predates the divergence of the Lemnaceae, we show that only *W. australiana* has also lost the MTI signaling components SOBIR1-BIR1 (Gao et al. 2009; Liebrand et al. 2013; Albert et al. 2015; van der Burgh et al. 2019). Previous work has shown RLP-SOBIR1 signaling depends on EDS1 and PAD4 in *A. thaliana* (Pruitt et al. 2021). Despite the loss of *SOBIR1, W. australiana* did not appear to have enhanced susceptibility to the bacterial pathogens tested when compared to other Lemnaceae. Given available resources, we focused on developing and characterizing the pathosystem of *S. polyrhiza* and *L. punctata* interactions with *Pst* DC3000 and *Pss* B728a. Transcriptional and bacterial mutant assays revealed the importance of phytohormones in the response of Lemnaceae species to pathogens; however, differential expression of typical marker gene orthologs was rarely observed. Furthermore, we show that MiAMP1 domain containing proteins are expanded in Lemnaceae and differentially expressed upon pathogen infection.

Duckweeds are known to exist with complex bacterial communities similar to those of the terrestrial leaf microbiome (Acosta et al. 2020), and a number of bacteria co-occurring with duckweeds have been shown to have growth promoting effects on duckweeds (O’Brien et al. 2020). To understand how the disease resistance of Lemnaceae to pathogenic bacteria is affected by the varying copy number of immune components, we screened a range of bacterial pathogens to identify a model pathosystem for further analysis. Our data demonstrate disease phenotypes upon inoculation with multiple bacterial pathogens. In addition, there were differences in macroscopic symptoms across Lemnaceae species upon treatment with the same pathogen. There was notable variability in the symptoms displayed by fronds within the same well with a given treatment, suggesting there may be environmental and population level factors that affect the outcome of infection. This variability in susceptibility across time points is consistent with the concept of quantitative resistance. The invasive nature of duckweeds and numerous healthy populations in the environment raises the question of how environmental factors in natural populations may contribute to duckweeds’ quantitative disease resistance. Furthermore, because duckweeds primarily reproduce clonally and have low mutation (Sandler et al. 2020) and effective recombination rates (Ho et al. 2019), there is limited opportunity for immune proteins to diversify, which has been shown to be important for generating new resistance specificities in other species. The rather unique combination of life-history traits of duckweeds including their fast-growth, clonal reproduction, reduced morphological complexity and aquatic habitat could theoretically all play a role in favoring quantitative means of resistance. For instance in a freshwater habitat the frequency of contact with PAMPs and microbes likely differs from soil and could therefore affect the balances between growth and defense; as energy spent on cell priming upon pathogen molecule perception reduces energy available for growth (Huot et al. 2014).

The reduced ability of *Pst* DC3000 hrcC^-^ to proliferate compared to wildtype suggests that the type-III secretion system and or the effectors it delivers can actively promote virulence in *S. polyrhiza* as well as in *A. thaliana* (Xin et al. 2016; Huot et al. 2017; Hauck, Thilmony, and He 2003). Furthermore, our results suggest that virulence of *Pst* DC3000 on *S. polyrhiza* relies on manipulation of host phytohormone pathways through the production of the JA mimic coronatine, similar to the interaction with *A. thaliana* (Moore et al. 1989; Mittal and Davis 1995). In contrast, *L. punctata* appears to have a stronger quantitative resistance response to *Pst* DC300 even in the presence of coronatine. Our data suggests that, despite the lack of conservation of the EDS1 pathway in *S. polyrhiza* and *L. punctata*, phytohormones and bacterial toxins remain key mediators of plant-bacterial infection. The reduction of MTI and ETI components in duckweeds opens up questions of how duckweed pathogens evolve; would reduction in effector repertoire be selected for due to a reduced pressure for MTI/ ETI suppression or would effectors further expand without an ETI system to keep them in check.

Consistent with the variation in phenotypes seen in bacterial inoculations, there was a lot of variation that was not explained by pathogen treatment between transcriptomes of duckweed biological replicates. The high variability could be a result of the asynchronous developmental stage of fronds, the quantitative nature of the resistance that we observed within the same population of duckweed or a combination of both of these factors. The use of whole plant tissues likely diluted the signal of differential gene expression. In the future, single cell transcriptomes might be more informative for interactions that involve quantitative resistance phenotypes. Despite the variability, our transcriptomic analysis revealed that the genes most consistently and differentially expressed after *Pseudomonas* treatment included MiAMP1 proteins. This suggests that the upregulation of MiAMP1 domain containing protein expression was consistent enough to be detected despite the overall variability. Such strong upregulation invites the speculation that they could provide alternative means of defense against invading pathogens similar to AMPs in Macadamia (McManus et al. 1999; Stephens et al. 2005) and Citrus (Huang et al. 2021). Other commonly differentially expressed protein families were associated with cytochrome P450s and phytohormones, indicating that specialized metabolites may have an important role in the infection response of *S. polyrhiza* and *L. punctata*. Together, our data highlights that, despite the absence of EDS1, there are conserved areas of immune response pathways, such as phytohormones and secondary metabolism. We also identified MiAMP1 proteins as strong candidate members of an alternative host immune response. Perhaps the reduced reliance on NLR - EDS1 pathways has released selective constraints allowing duckweeds to avoid the typical land plant NLR - pathogen effector arms races that are burdening other plant species and facilitated amplification of alternate immune strategies.

Duckweeds are an exciting system for future research into the plant immune system. They provide a reduced redundancy system in terms of both gene copy number and pathways present. In addition, their rapid lifecycle of just 34 hrs (Michael et al. 2020), small size and susceptibility to model pathogens make them the perfect system for high-throughput experimentation. The genomic and genetic resources are also developing at a rapid pace, with multiple genomes available (Michael et al. 2017; An et al. 2019; Harkess et al. 2021; Ho et al. 2019; P. N. T. Hoang et al. 2018; W. Wang et al. 2014; S. Xu et al. 2019; Michael et al. 2020) and a range of transformation protocols (Chanroj, Jaiprasert, and Issaro 2021; Acosta et al. 2021; Liu et al. 2019; J. Yang et al. 2018; G.-L. Yang et al. 2018; Ko et al. 2011). Many interesting questions remain to be answered including how duckweed plants are able to thrive in a wide range of environments despite being highly susceptible to bacterial phytopathogens in the laboratory setting. A hypothesis is that duckweeds may be using means of disease protection that are inherently absent in the laboratory due to the way duckweeds are propagated. This may include microbiome-mediated disease protection, small peptide defense molecules such as MiAMP1s or chemical defense strategies. A combination of genetics, molecular biology and metabolomic approaches will be needed to address such hypotheses.

## Materials and Methods

### *Landoltia punctata* clone 5635 DNA isolation and sequencing

*L. punctata* clone 5635 (Lp5635, formerly DWC138) was received from the Rutgers Duckweed Stock Collective (RDSC; http://www.ruduckweed.org/). Lp5635 was collected, patted dry to remove excess water and flash frozen in liquid nitrogen. Frozen tissue was ground in a mortar and pestle under liquid nitrogen. High Molecular Weight (HMW) DNA was isolated using a modified Bomb protocol (https://doi.org/10.1371/journal.pbio.3000107). DNA quality was assessed on a Bioanalyzer and HMW status was confirmed on an agarose gel. Libraries were prepared from HMW DNA using NEBnext (NEB, Beverly, MA) and 2×150 bp paired end reads were generated on the Illumina NovaSeq (San Diego, CA). Resulting raw sequence was only trimmed for adaptors, resulting > 60x coverage of the diploid *L. punctata* genome (350 Mb).

### *Landoltia punctata* clone 5635 genome assembly, gene prediction and annotation

Illumina paired end reads were assembled with Spades (v3.14.0) with the default settings (Bankevich et al. 2012). Resulting contigs were annotated using a pipeline consisting of four major steps: repeat masking, transcript assembly, gene model prediction, and functional annotation as described (Abramson et al. 2021). Repeats were identified using EDTA (v1.9.8) (Ou et al. 2019) and these repeats were used for softmasking. RNAseq reads were aligned to the genomes using minimap2 and assembled into transcript models using Stringtie (v1.3.6). The soft-masked genome and Stringtie models were then processed by Funannotate (v1.6) (https://github.com/nextgenusfs/funannotate) to produce gene models. Predicted proteins were then functionally annotated using Eggnog-mapper (v2) (Huerta-Cepas et al. 2017).

### Pathway ortholog identification

To estimate the date of the loss of *EDS1, PAD4, ADR1* and *NDR1*, we looked for species closely related to the Lemnaceae with available genome sequence. We then ran a BLASTP search using the *Oryza sativa* and *Arabidopsis thaliana* EDS1, PAD4, ADR1 and NDR1 proteins as queries against the Taro genome (https://doi.org/10.1111/1755-0998.13239). To check orthology, the subject sequence identified in the Taro proteome was then blasted back to the *Oryza sativa* and *Arabidopsis thaliana* proteomes using the Phytozome BLAST tool. Upon confirming the presence of EDS1, PAD4, ADR1 and NDR1 in Taro, we then utilized TimeTree.org to estimate the time window during which EDS1, PAD4, ADR1 and NDR1 were lost.

To identify orthologous gene groups, Orthofinder (v2.5.4) (Emms and Kelly 2019) was used on the proteomes of *Spirodela polyrhiza* 7498 (W. Wang et al. 2014), *Landoltia punctata* 5635, *Wolffia australiana* 8730 (Michael et al. 2020) and *A. thaliana* TAIR 10 (Lamesch et al. 2012). Pathway gene clusters were identified from a literature search of *A. thaliana* effector triggered immune responses and EDS1/PAD4 interacting proteins (S.table 8). These Arabidopsis proteins were then used to pull out corresponding orthogroups containing Lemnaceae species which were cross checked using Phytozome for presence in *S. polyrhiza*. For instances of absence, gene loss was verified using tBLASTn (Camacho et al. 2009). For large gene family analysis of MiAMP1, RLK, RLP and NLRs, pfamscan (Madeira et al. 2019; Sarris et al. 2016) was used to identify proteins with domains of interest (e-value = 10; MiAMP1, RLK, RLP and NB-ARC domains) (S.table1). RLK, RLP and MiAMP1 protein sequences were aligned with MUSCLE v3.8.1551 (Madeira et al. 2019) and NB-ARC domains were aligned with hmmalign from HMMER3 (Wheeler and Eddy 2013) to NB-ARC1-ARC2 (Bailey et al. 2018). Alignments were manually curated using BELVU (Barson and Griffiths 2016) and Jalview (Waterhouse et al. 2009). Maximum likelihood phylogenies were calculated utilizing MPI version of RAxML (v8.2.9) (-f a, -x 12345, -p 12345, -# 100, -m PROTCATJTT).

### Plant growth conditions

Duckweed fronds were propagated by transferring 3 mother fronds to a new well containing fresh media at a frequency of 3-4 weeks. Plants were grown on Schenk and Hildebrandt basal salt media (Sigma-Aldrich, S6765-10L) (0.8% agarose, pH 6.5) in 6- or 12-well plates and then placed in a growth chamber set to 23°C with a diurnal cycle of 16 hr light (75 μmol) / 8 hr dark.

### Pathogen inoculation

Bacterial colonies were grown in Luria Broth (LB) media supplemented with the following antibiotics: [10 μg/ml; kanamycin (Km), 50 μg/ml; rifampicin (Rif), 50 μg/ml; spectinomycin (Sp)] overnight at 28°C in a shaking incubator (20.5 g). Pathogens used in this study included: *Pseudomonas syringae* pv. tomato DC3000 GFP (Matthysse et al. 1996; Mudgett and Staskawicz 1999), *P. syringae* pv. tomato DC3000 cma^-^ (Sreedharan et al. 2006), *P. syringae* pv. tomato DC3000 hrcC^-^ (Mudgett and Staskawicz 1999), *P. fluorescens* N2C3 (DSM 106121) (Parte et al. 2020), *P. syringae* B7281 (Feil et al. 2005), *P. syringae* pv. glycinea race 4 (Staskawicz et al. 1987), *P. syringae* pv tabaci 11528 (Institute of Medicine (US) Committee on Resource Sharing et al. 1996), P. syringae pv. maculicola (Davis, Schott, and Ausubel, n.d.), *Xanthomonas euvesicatoria* (Roden et al. 2004), X. gardneri (Schwartz et al. 2017), *X. perforans* (Bophela et al. 2019), and *X. translucens* (Peng et al. 2016). All strains were plated on Rif; *P. syringae pv. tomato* DC3000 on Rif/Km and *P. syringae pv. tomato DC3000* Δcmfa on Rif/Km/Sp. Liquid culture was spun down on a benchtop centrifuge at 3,000 g for 15 minutes. The pellet was then resuspended in 10 mM MgCl_2_, OD_600_ was determined and the infiltration solution was diluted to a final standard high inoculum of OD_600_= 0.1, equivalent to 1×10^8^ cells of *Pst* DC3000. A total of 500 μL of OD_600_= 0.1 bacterial inoculation solution was pipetted on to 3 duckweed fronds per well. The plate was then either placed straight into the incubator or placed under vacuum (0.8 PSI) for 10 minutes.

For growth curves, bacterial cells were resuspended at 1×10^8^ CFU/ml, OD_600_=0.1 in 10 mM MgCl_2_. The inoculum was further diluted to a standard low inoculum with a final concentration of 1×10^5^CFU/ml. Each well was then inoculated with 500 μl of treatment, followed by vacuum infiltration (0.8 PSI) for 10 minutes. For each timepoint 1 cm^2^ of fronds was sampled in 150 μl MgCl_2_ and glass beads, and homogenized in a biospec mini-beadbeater at 2000 rpm with vital distance 1.25 inch. Serial dilutions were made and plated on appropriate selective media. Two days after plating, colonies were counted.

### Microscopy

For microscopy, flood inoculation with 500 μL solutions OD_600_=0.1 were used. Whole fronds were staged in water on slides and covered with a glass coverslip. The slides were imaged on a Zeiss 710 LSM confocal microscope with either the 20x (water), 63x (oil) or 100x (oil) objectives. To image bacteria on duckweed fronds, they were stained with 10μl of 1x Syto™ BC Green Fluorescent Nucleic Acid Stain (Thermo Scientific, S34855).

### Hormone supplementation

Coronatine (Sigma-Aldrich, C8115-1MG) powder was dissolved in 100% DMSO to create a 200 μM/ml stock that was then diluted to 3 μM and 0.3 μM in ddH_2_O. Salicylic acid BioXtra ≥99.0% (Sigma-Aldrich, S5922-100G) was diluted in water to concentrations of 2mM, 0.4mM and 0.2mM. Buffer solution used was the same as the solvent for the treatment. Individual wells of 6 or 12 well plates containing 3 duckweed fronds were inoculated with 500 μL or 250 μL of phytohormone solution, respectively. Phytohormone treatments were applied just prior to pathogen treatment.

### RNAseq analysis

Inoculations were conducted as outlined above using standard high bacterial inoculum 1×10^8^CFU/ml. At 30 minutes, 1 hr, 6 hrs and 12 hrs after inoculation, fronds from the same well and the same treatment were transferred to microcentrifuge tubes containing 1.5 ml RNAlater and frozen in liquid nitrogen. RNA was extracted using Qiagen RNAeasy plant kit (74903) and only samples with a RIN score of >8.0 from Agilent Bioanalyzer 2100 performed by QB3-UC Berkeley were used for library preparation. Library construction and sequencing were performed by Novogene using NEBNext Ultra II RNA library prep by Illumina (E7770) and an Illumina Novaseq 6000 S4 instrument. After sequencing, we demultiplexed reads using Illumina indices and then ran a QC check involving filtering N containing, low quality, and adapter-related reads. Upon receiving the data from Novogene, an initial quality check of reads was performed with fastqc (“Babraham Bioinformatics - FastQC A Quality Control Tool for High Throughput Sequence Data” n.d.). Reads were then mapped to the *S. polyrhiza* v2 genome using HISAT2 (Kim, Langmead, and Salzberg 2015). Read coverage tables were computed using stringtie (Pertea et al. 2015) and differential gene expression analysis was carried out using EdgeR (Dai et al. 2014). Genes were considered differentially expressed if they met the criteria of |Log_2_FC| >1, FDR < 0.05. For a detailed list of commands see: https://github.com/erin-baggs/DuckweedRNA. Outliers were removed by visual inspection; the edgeR count tables were plotted as PlotMDS method Log2FC and BCV (Sfig 16-19). Samples were removed if they alone were causing most of the variance on a dimension leading to all other samples clustered into a corner. Then, if one sample of a treatment was grouping with samples of an opposing treatment, marker gene expression was analysed to check whether the sample’s expression pattern may have been the result of cross contamination. If the expression was inconsistent with at least 3 other replicates from the treatment group it belonged to, the sample was removed. Replicates removed from *S. polyrhiza* analysis included B3, B20, B19, P21, B34, P114. The replicates B15, B20, P39, P70, H80, B94, P103 and H115 were removed from *L. punctata* analysis.

## Data availability

Sequencing reads for *S. polyrhiza* and *L. punctata* RNAseq can be found at the respective NCBI BioProjects PRJNA808038 and PRJNA801691. The *L. punctata* 5635 genome can be found on CoGe accession ID: 63585. Scripts for analysis presented in the manuscript can be found on the github repository https://github.com/erin-baggs/DuckweedRNA. Z-stack of *Pst* DC3000 infection of *S. polyrhiza* is available from Zenodo doi: 10.5281/zenodo.5639580.

## Acknowledgements

We would like to thank Dr. Denise Schichnes and Dr. Steven Ruzin for their expert advice and help with confocal microscopy. Many thanks to China Lunde for propagation of duckweed during the pandemic and to Dr. Wilfied Haerty and the members of the Krasileva lab for helpful discussions. We acknowledge funding from the Biotechnology and Biological Sciences Research Council (BBSRC), part of UK Research and Innovation, Core Capability Grant BB/CCG1720/1 and the work delivered via the Scientific Computing group, as well as support for the physical HPC infrastructure and data centre delivered via the NBI Computing infrastructure for Science (CiS) group. KVK has been funded by the Gordon and Betty Moore Foundation, Innovative Genomics Institute, and the NIH Director’s Award.

## Author contributions

E.L.B. performed genomic/transcriptomic presence absence analysis, growth curves, hormone assays, microscopy and RNAseq analysis. M.B.T and E.L.B. performed the screen of phytopathogens across duckweed species and RNA extractions. B.W.A and T.P.M performed DNA extraction, genome assembly and annotation of *Landoltia punctata* DWC138 genome. E.L.B and K.V.K designed the study. E.L.B prepared figures and wrote the full manuscript draft; all authors have read and approved the final version of the manuscript.

## Supplemental datasets

### Supplemental tables

S.table 1 - Copy number of orthologs/homologs of immune associated genes.

S.table 2 - Table of Spirodela polyrhiza homologs to Arabidopsis immunity genes.

S.table 3 - Syndrome prevalence upon pathogen treatment of Lemnaceae species.

S.table 4 - Log2FC of JAZ domain containing *S. polyrhiza* genes.

S.table 5 - Log2FC of WRKY domain containing *S. polyrhiza* genes.

S.table 6 - Log2FC of MiAMP1 domain containing *S. polyrhiza* genes.

S.table 7 - Log2FC of NB-ARC domain containing *S. polyrhiza* genes.

S.table 8 - Log2FC of JAZ domain containing *L. punctata* genes.

S.table 9 - Log2FC of WRKY domain containing *L. punctata* genes.

S.table 10 - Log2FC NBARC domain containing genes *L. punctata*.

S.table 11- Log2FC MiAMP1 domain containing genes *L. punctata*.

S.table 12 - Identification of homologs for transcriptomic analysis between *A. thaliana, S. polyrhiza* and *L. punctata*.

S.table 13 - Orthologs to *A. thaliana* marker genes that are significantly upregulated upon *Pst* treatment Log2FC >0.75 FDR<0.05 in either *S. polyrhiza* or *L. punctata*.

### Supplemental datasets

Supplemental datasets are available at github (https://github.com/krasileva-group/Thesis-Scripts) or zenodo.

Sdataset 1 - Powerpoint presentation of pathogen inoculation symptoms - doi: 10.5281/zenodo.5733456

Sdataset 2 - Zstack microscopy - doi: 10.5281/zenodo.5639580

Sdataset 3 - Differentially expressed gene tables *S. polyrhiza* - github

Sdataset 4 - Differentially expressed gene tables *L. punctata* - github

Sdataset 5 - Orthogroup assignment and overlap across pathogen treatments and species - github

