## Supplemental Figures for "Characterization of EDS1-independent plant defense responses against bacterial pathogens using Duckweed/*Pseudomonas* pathosystems"

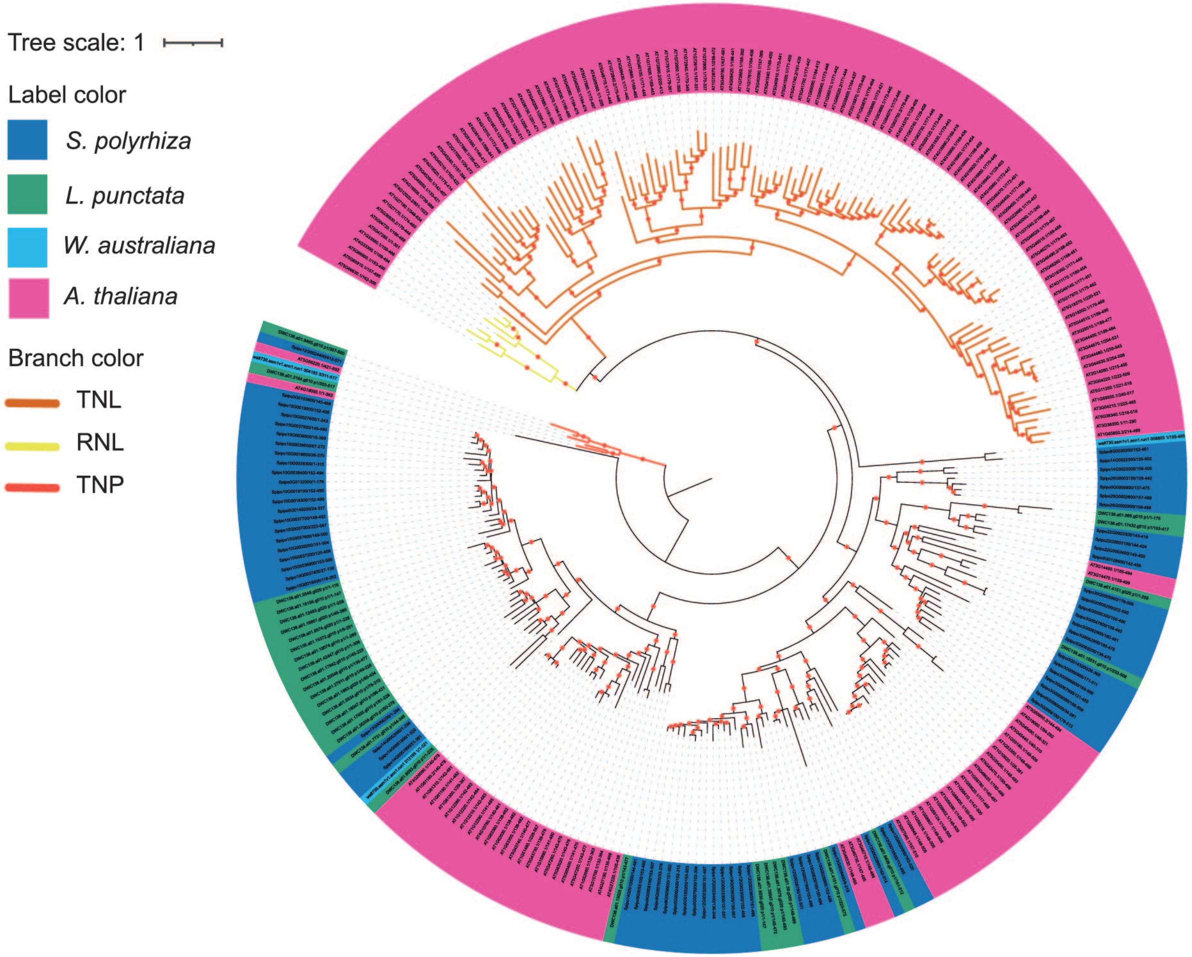


**Sfigure 1. Phylogeny of NLR proteins in duckweeds.**
Maximum likelihood phylogeny of NB-ARC proteins (Pfam PF00931) from *A. thaliana, S. polyrhiza*, *L. punctata* and *W. australiana*. Alignments were manually curated for the presence of NB-ARC functional motifs. Red dots indicate bootstrap values >70. Orange branches indicate TIR-NBARC-LRR proteins, red branches indicate TIR-NBARC-like-TPR repeat containing proteins and yellow branches indicate RPW8-NB-ARC-LRR (RNLS). Gene identifiers are colored to indicate species: *S. polyrhiza* - dark blue, *W. australiana* - light blue, *L. punctata* - green and *A. thaliana* - pink. Tree rooted on TNPs.

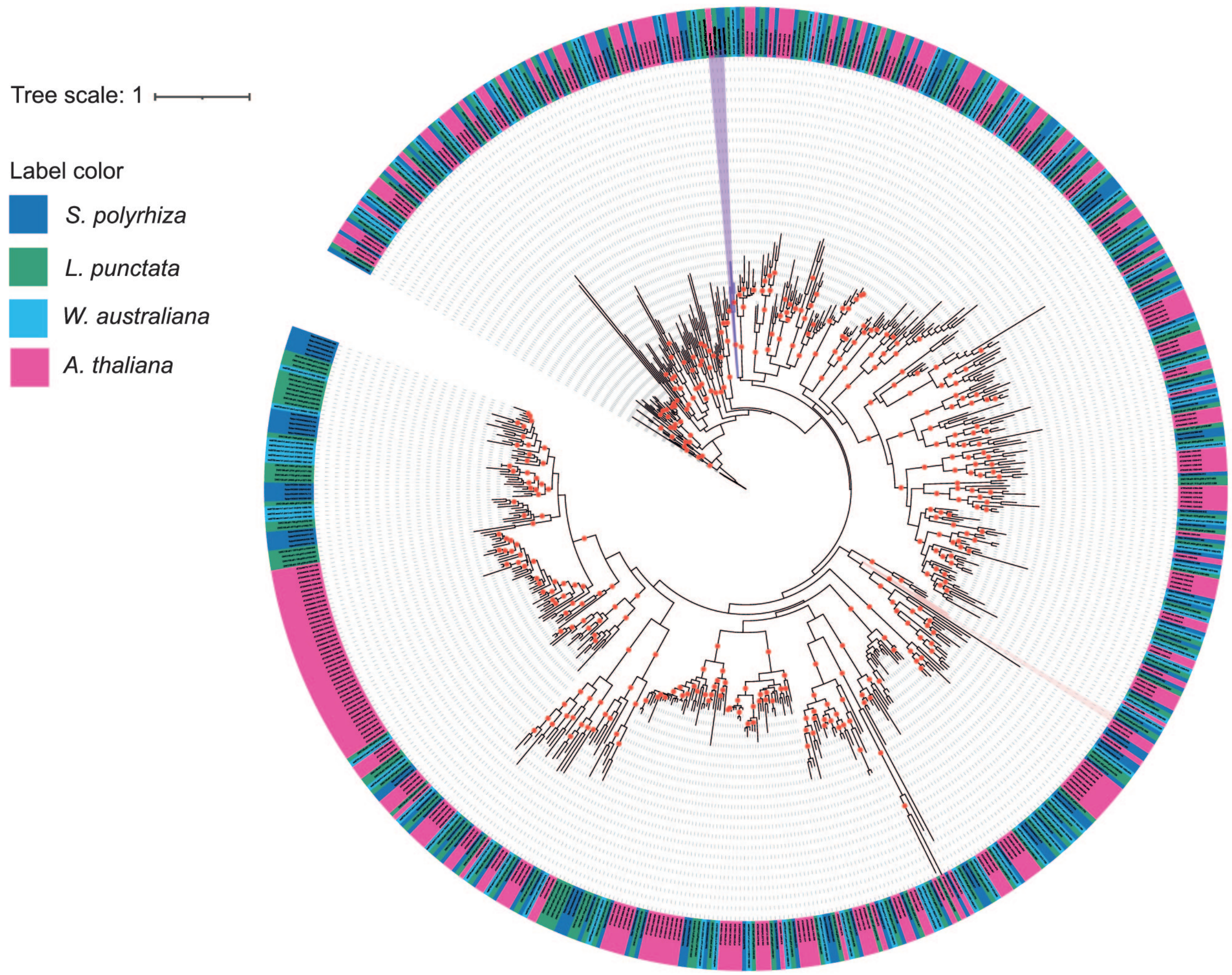


**Sfigure 2. Phylogeny of RLK proteins in duckweeds.**
Maximum likelihood phylogeny of RLK proteins from *A. thaliana, S. polyrhiza, L. punctata* and *W. australiana*. Alignments were manually curated for the presence of RLK functional motifs. Red dots indicate bootstrap values >70. Tree rooted at midpoint. Gene identifiers are colored to indicate species: *S. polyrhiza* - dark blue, *W. australiana* - light blue, *L. punctata* - green and *A. thaliana* - pink. Purple range indicates SOBIR1 clade and pink range indicates FLS2 clade.

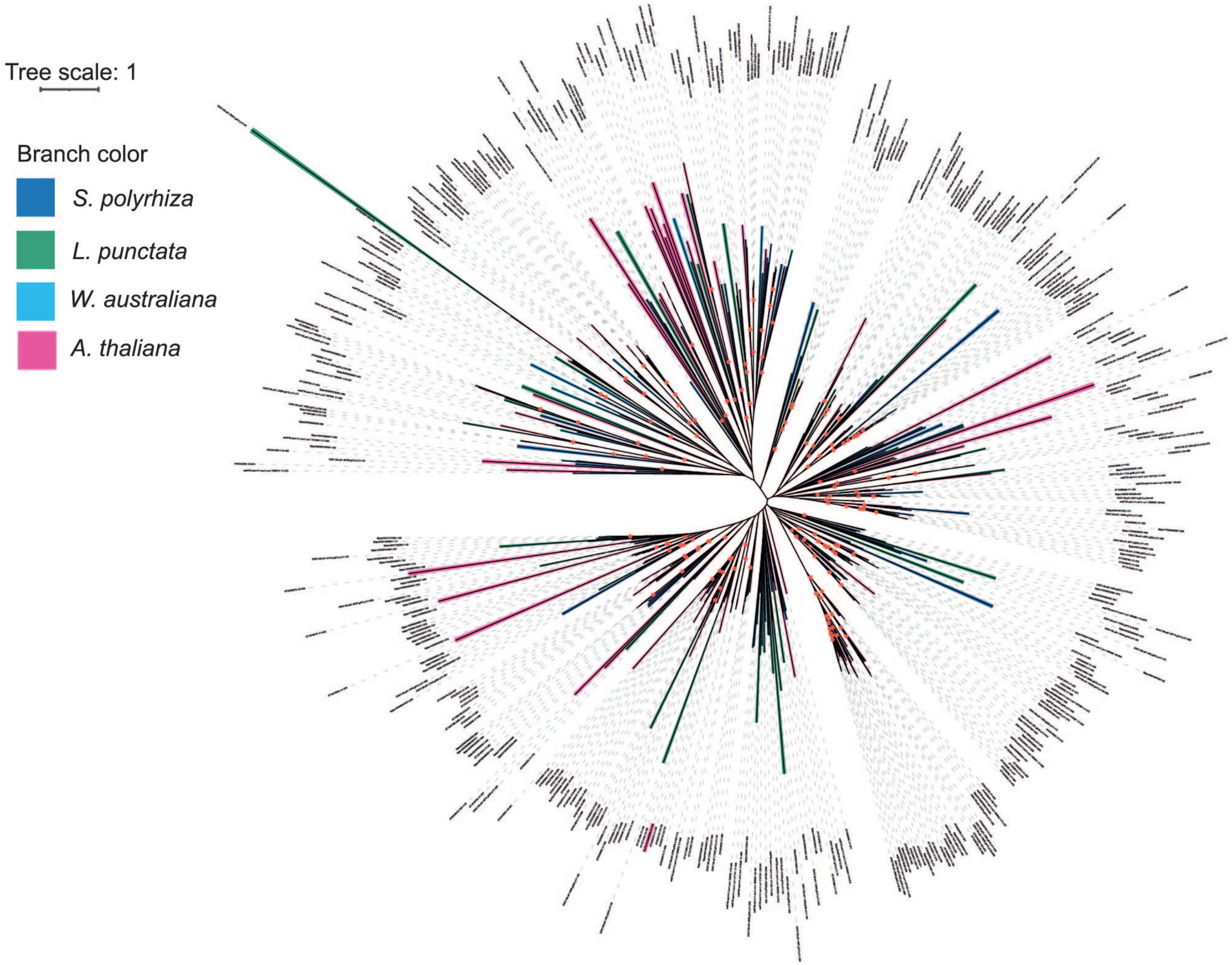


**Sfigure 3. Phylogeny of RLP-type proteins in duckweeds.**
Maximum likelihood phylogeny of RLP proteins from *A. thaliana, S. polyrhiza, L. punctata* and *W. australiana*. Alignments were manually curated for conserved motifs. Red dots indicate bootstrap values >70. Branches are colored to indicate species: *S. polyrhiza* - dark blue, *W. australiana* - light blue, *L. punctata* - green and *A. thaliana* - pink.

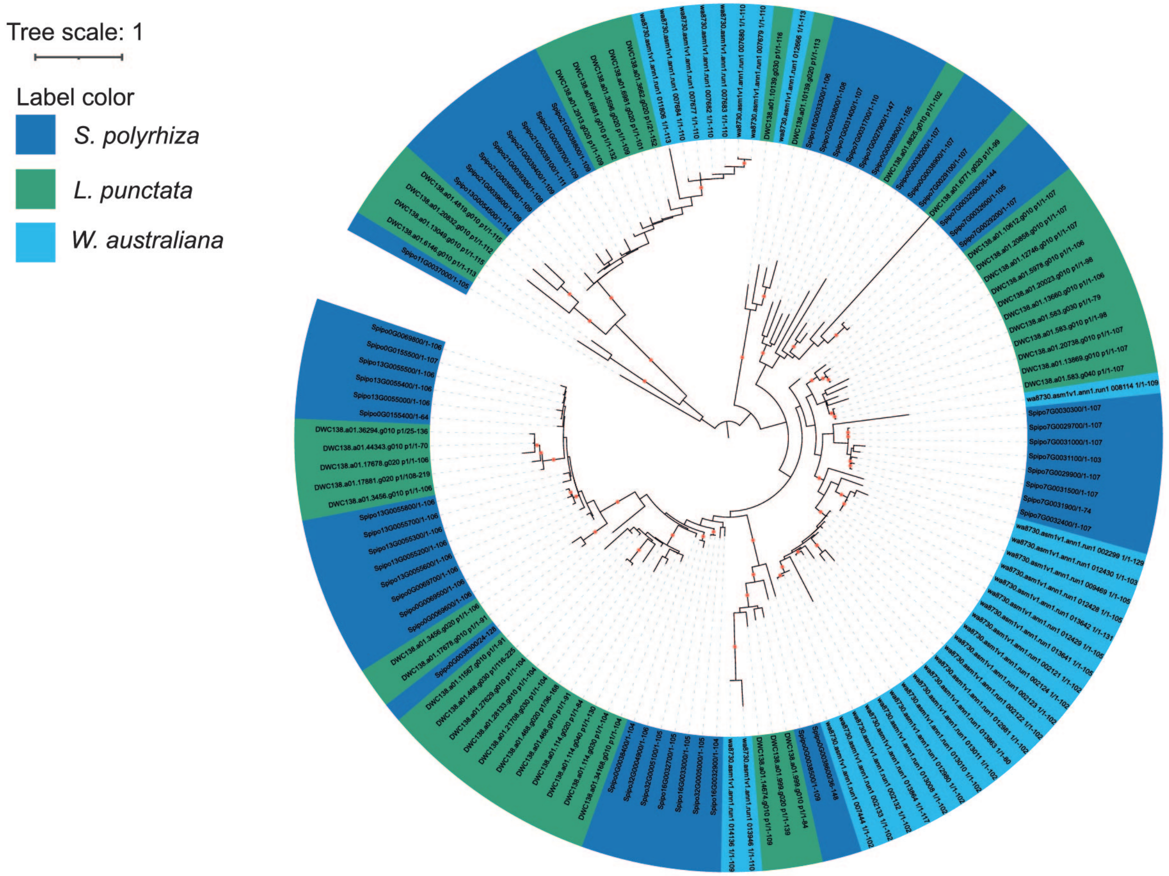


**Sfigure 4. Phylogeny of MiAMP1 domain containing proteins in duckweed species.**
Maximum likelihood phylogeny of MiAMP1 (Pfam PF09117 ) domain containing proteins from *A. thaliana, S. polyrhiza, L. punctata* and *W. australiana*. Red dots indicate bootstrap values >70. Tree rooted at midpoint. Gene identifiers are colored to indicate species: *S. polyrhiza* - dark blue, *W. australiana* - light blue and *L. punctata* - green.

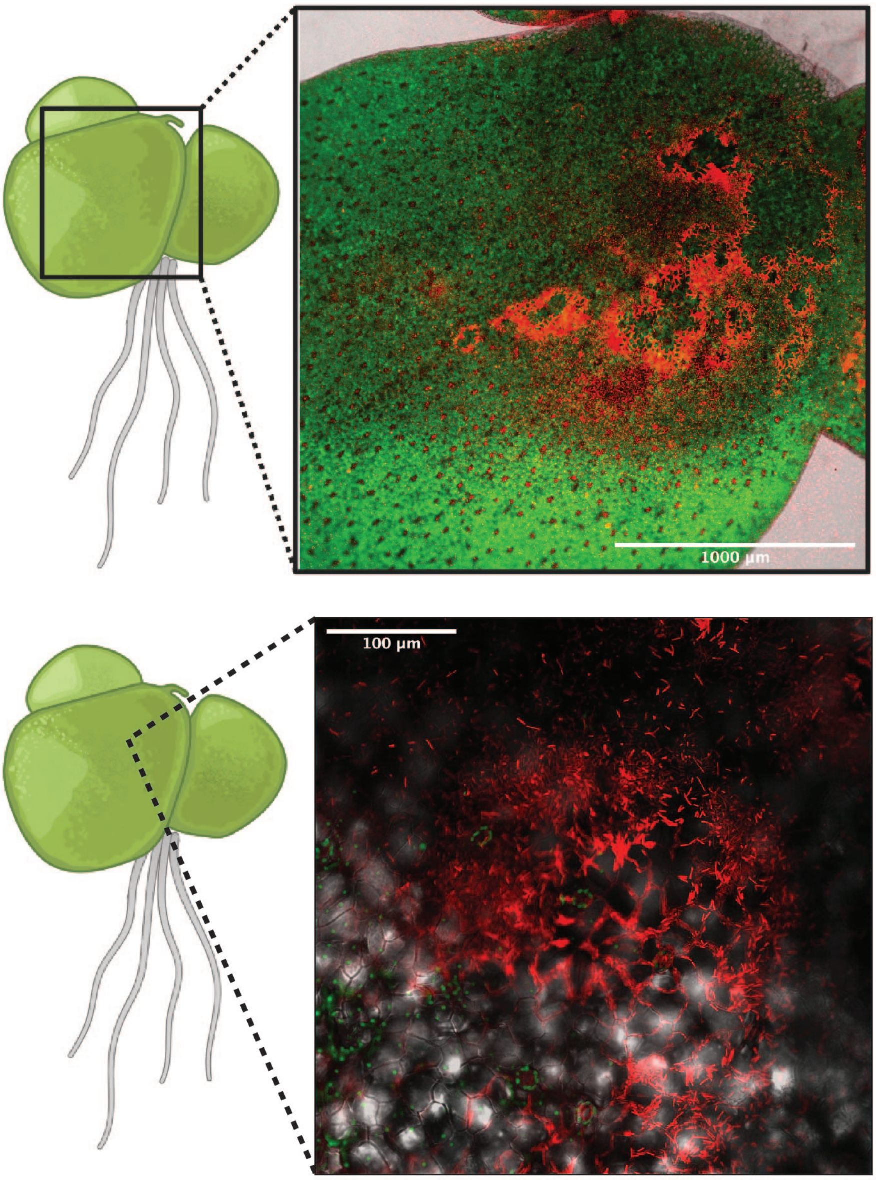


**Sfigure 5 Microscopy of frond surface 5 days post flood innoculation with *Pseudomonas syringae* pv. tomato DC3000**.
A. Confocal microscopy 5x magnification red false coloring represents *Pst* DC3000 stained with SytoBC, green shows chlorophyll fluorescence and grey is transmitted light. B. 20x confocal microscopy of concentrated Pst DC3000 populations.

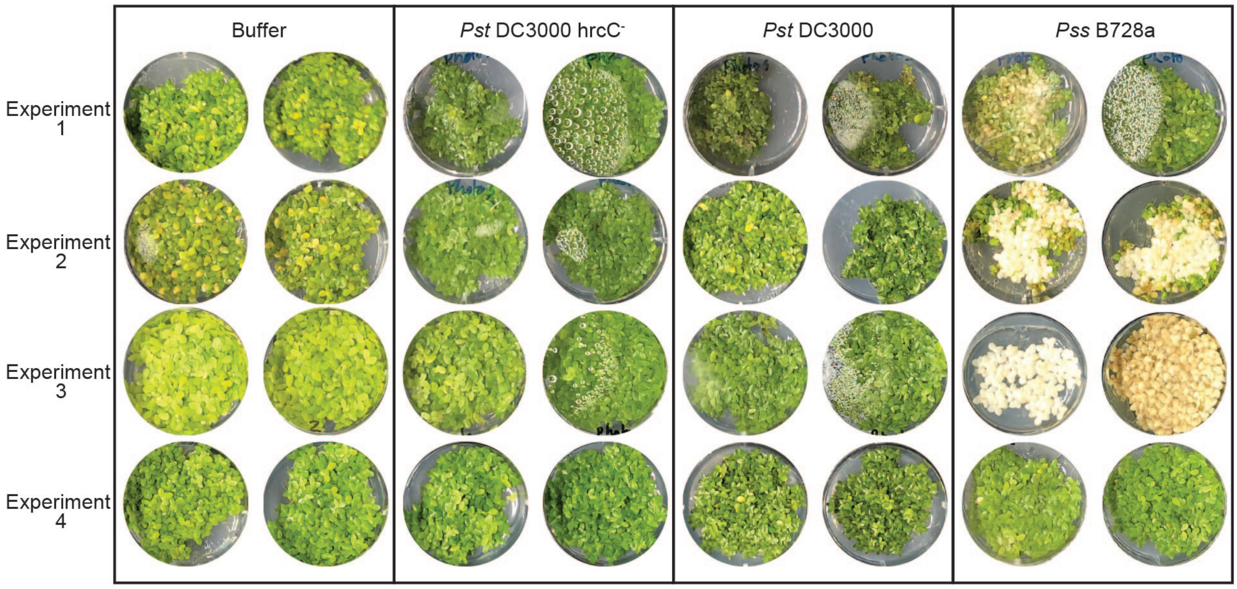


**Sfigure 6  Low bacterial load infection of Landoltia punctata 1 month post inoculation.**
All wells of an experiment started with 12 fronds and were treated on the same day, each well is a separate biological replicate. There are two replicates photographed per treatment. Some wells have condensation which can be seen as the water droplets obscuring fronds below, this is as plates need to be sealed to prevent contamination.

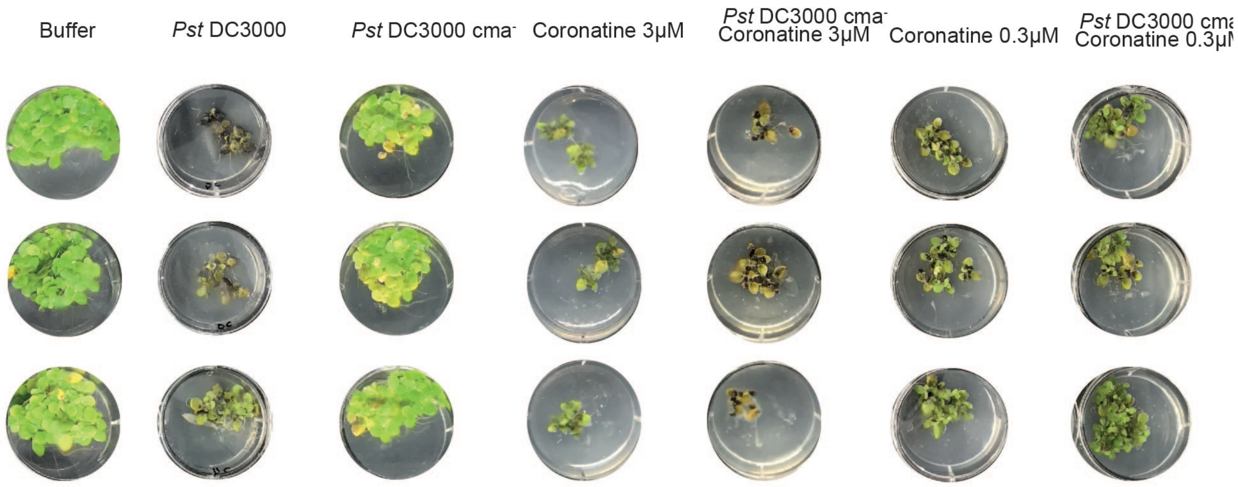


**Sfigure 7. Role of coronatine in *Pst* DC3000 infection of *Spirodela polyrhiza*.**
Experiment 2 of coronatine response experiment figure 4a. All wells were treated on the same day and each well is a separate biological replicate.

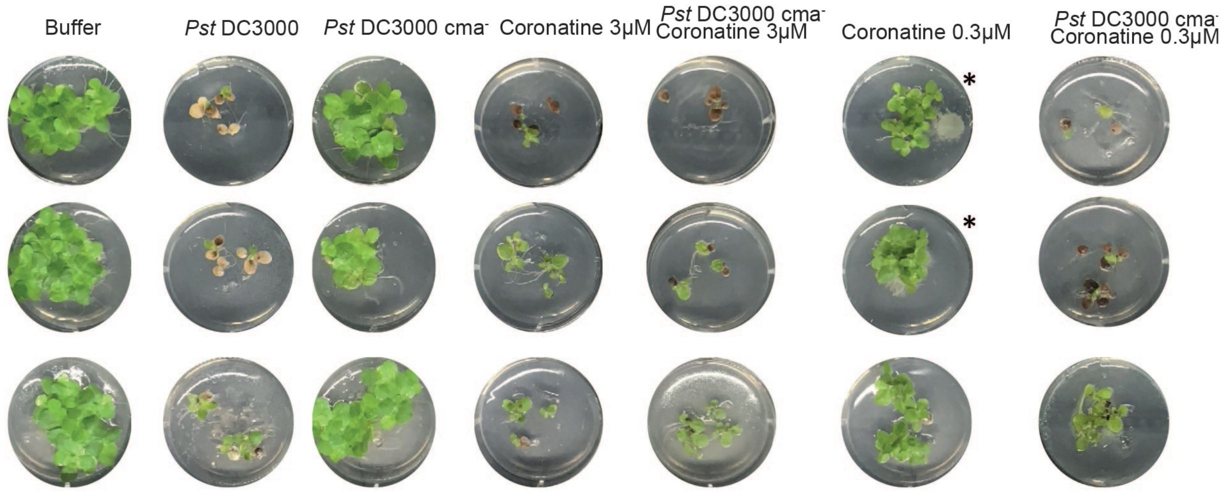


**Sfigure 8. Role of coronatine in *Pst* DC3000 infection of *Spirodela polyrhiza*.**
Experiment 3 of coronatine response experiment figure 4a. All wells were treated on the same day and each well is a separate biological replicate. Asterix marks wells with visible contamination.

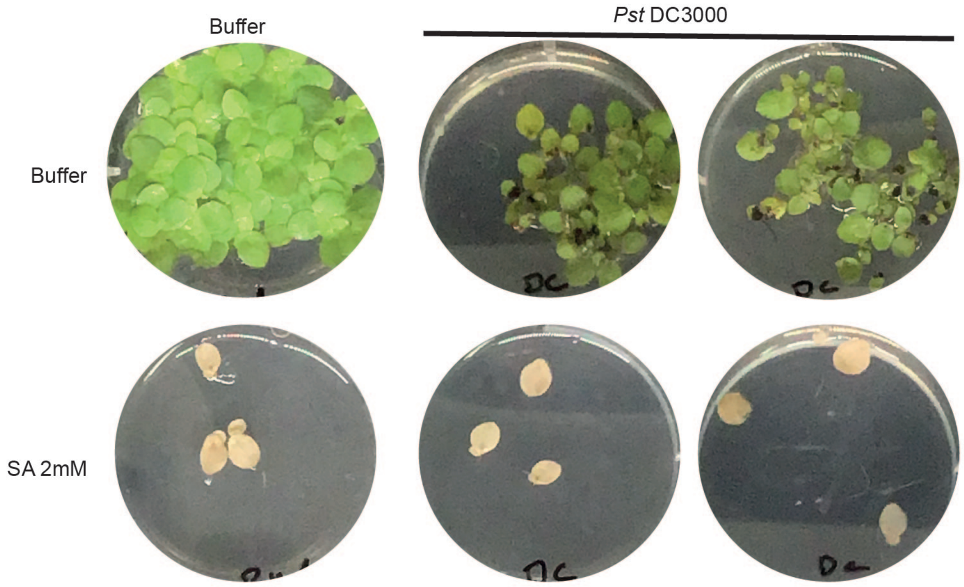


**Sfigure 9. Salicylic acid phytotoxicity to *Spirodela polyrhiza* upon buffer or *Pst* DC3000 treatment.**
Inoculation with SA 24hrs before pathogen inoculation. Images 3 weeks after treatment with buffer (10mM MgCl2) and *Pst* DC3000.

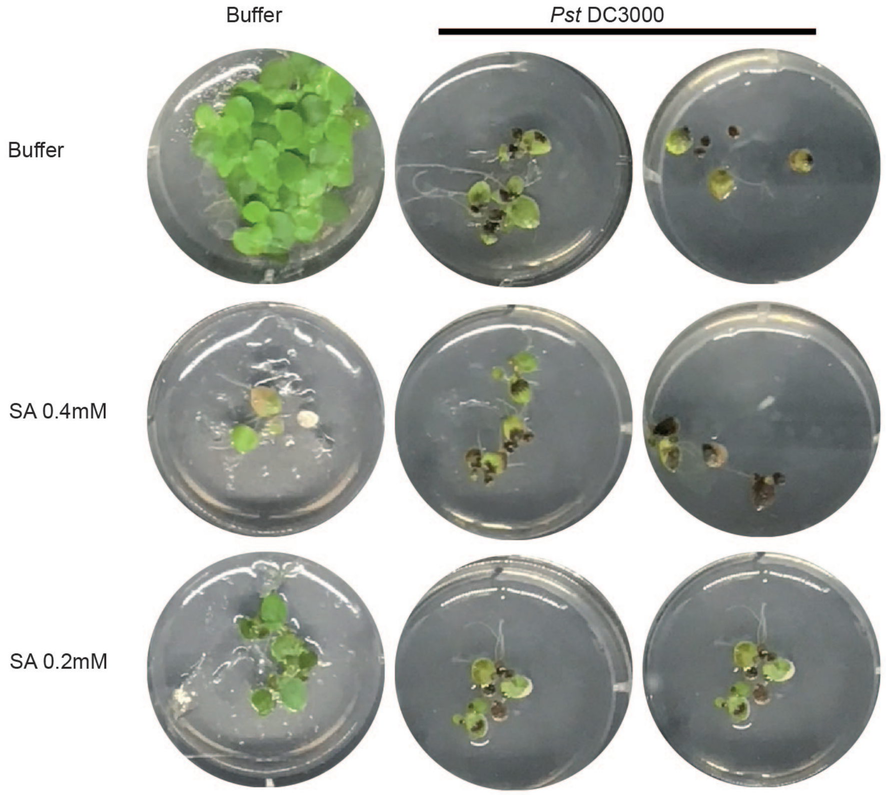


**Sfigure 10. Role of salicylic acid in *Pst* DC3000 infection of *Spirodela polyrhiza*.**
Experiment 2 of salicylic acid treatment of *S. polyrhiza* experiment figure 4b. All wells were treated on the same day and each well is a separate biological replicate.

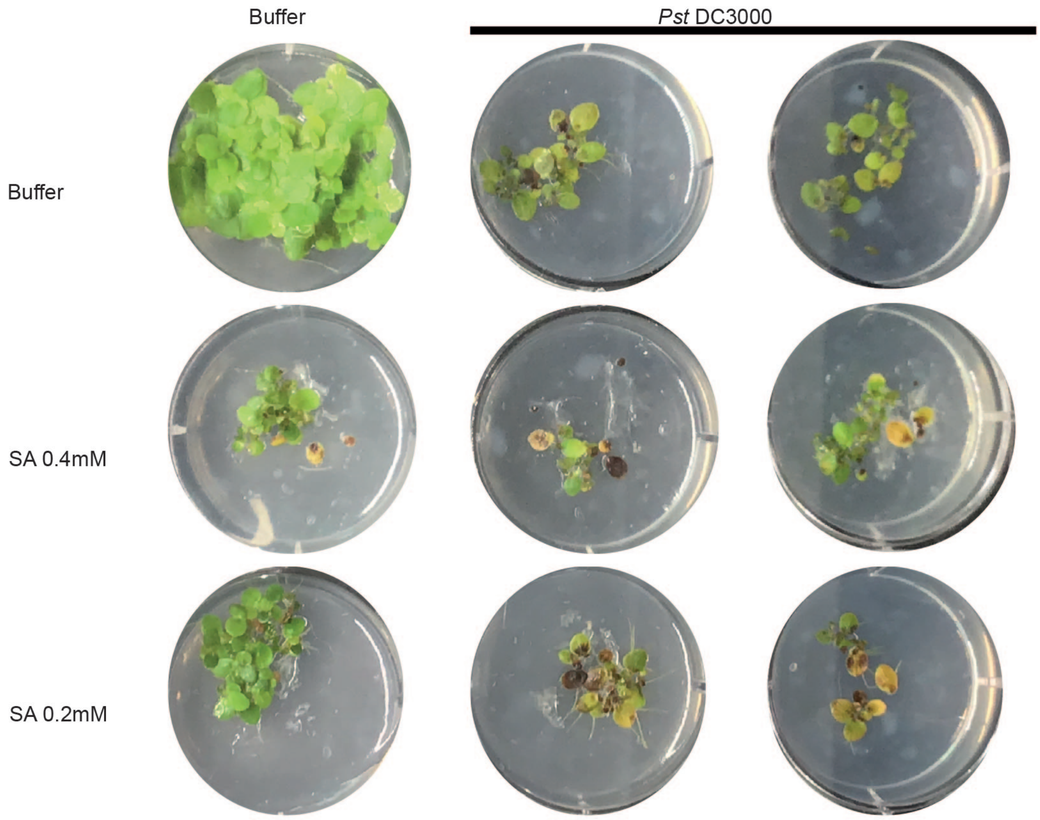


**Sfigure 11. Role of salicylic acid in *Pst* DC3000 infection of *Spirodela polyrhiza.***
Experiment 3 of salicylic acid treatment of *S. polyrhiza* experiment figure 4b. All wells were treated on the same day and each well is a separate biological replicate.

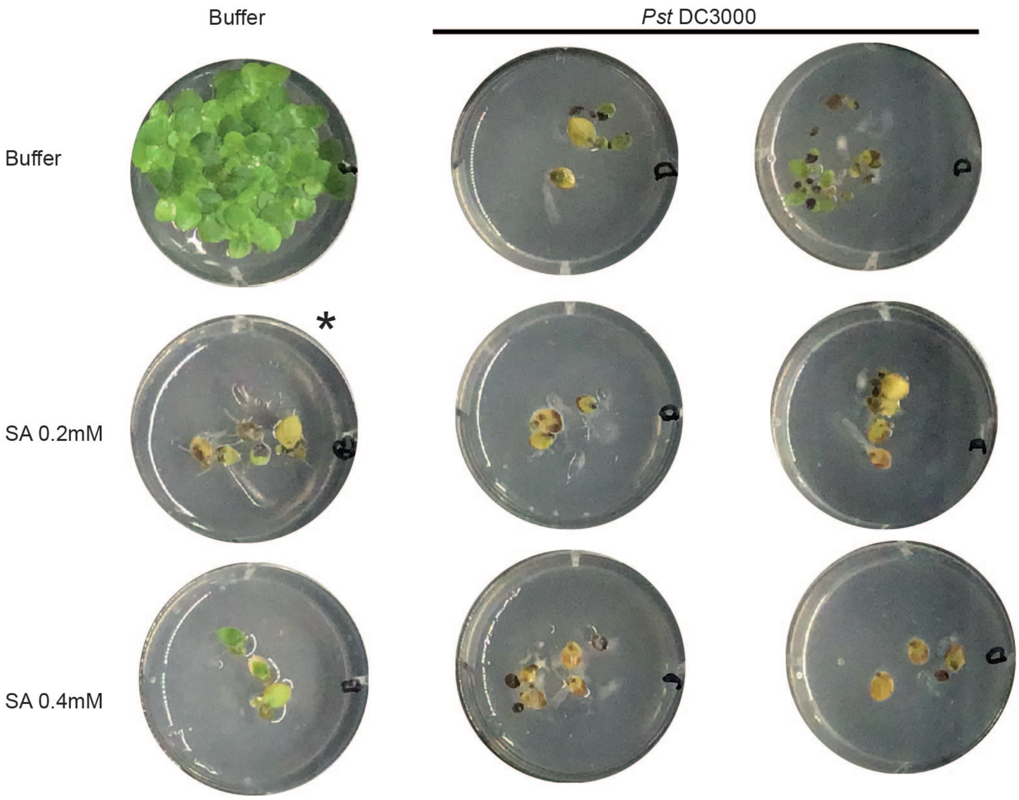


**Sfigure 12. Role of salicylic acid in *Pst* DC3000 infection of *Spirodela polyrhiza.***
Experiment 4 of salicylic acid treatment of *S. polyrhiza* experiment figure 4b. All wells were treated on the same day and each well is a separate biological replicate. Asterisk marks wellswith visible contamination of the buffer with *Pst* DC3000.

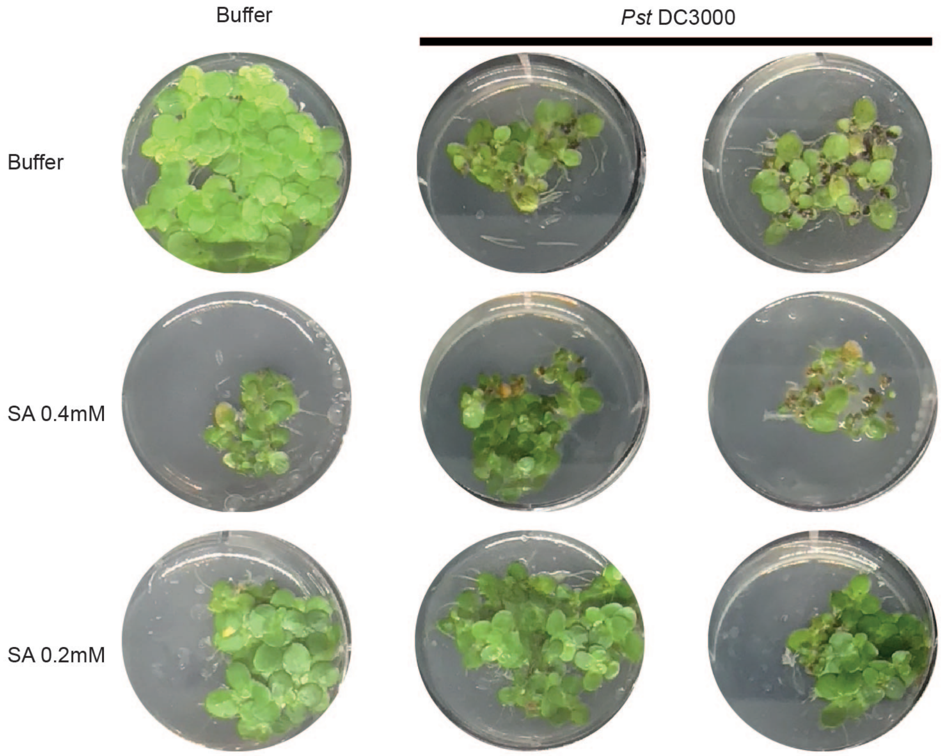


**Sfigure 13. Role of salicylic acid in *Pst* DC3000 infection of *Spirodela polyrhiza*.**
Experiment 5 of salicylic acid treatment of *S. polyrhiza* experiment figure 4b. All wells were treated on the same day and each well is a separate biological replicate. Asterisk marks wells with visible contamination of the buffer with *Pst* DC3000. .

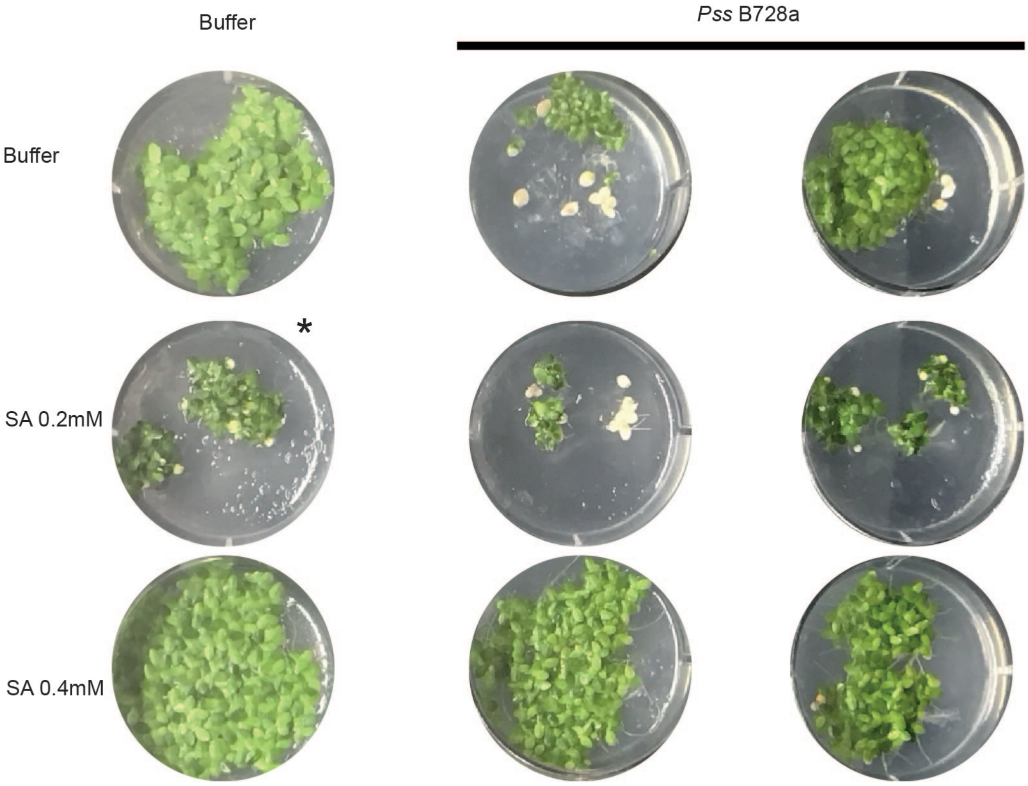


**Sfigure 14. Role of salicylic acid in *Pss* B728a infection of *Landoltia punctata*.**
Experiment 2 of salicylic acid treatment of *L. punctata* experiment figure 4c. All wells were treated on the same day and each well is a separate biological replicate. Asterisk marks wells with visible contamination of the buffer with *Pss* B728a.

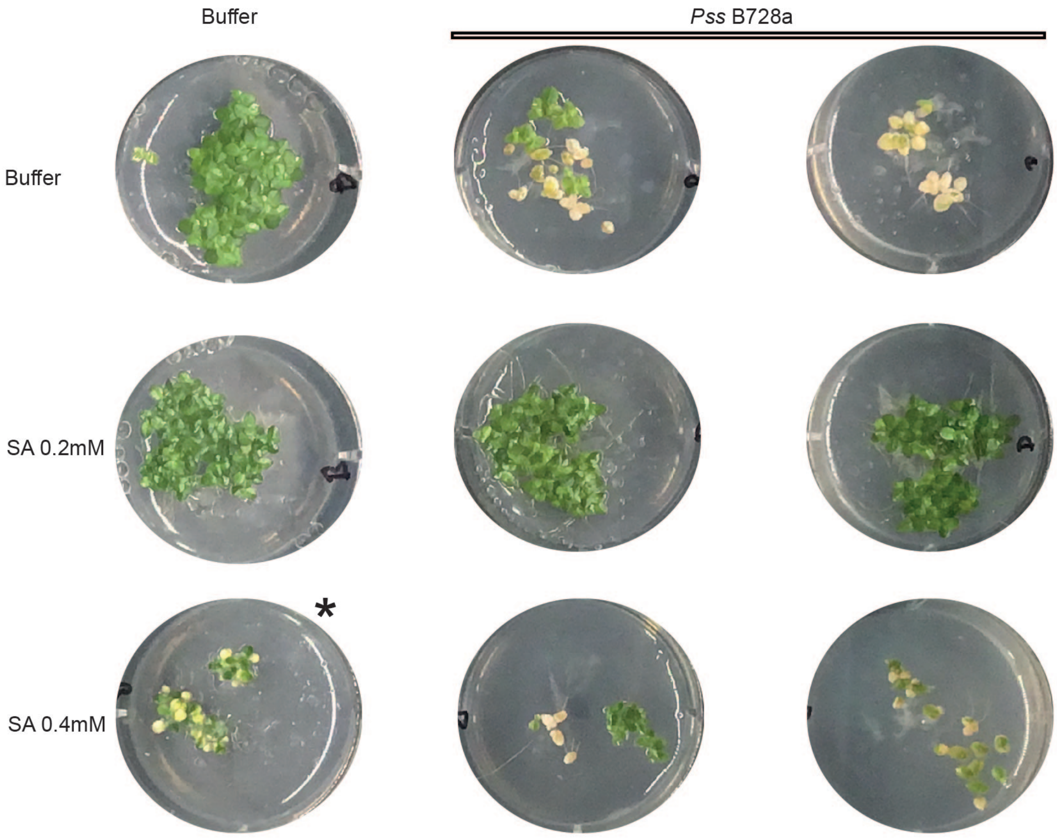


**Sfigure 15. Role of salicylic acid in *Pss* B728a infection of *Landoltia punctata.***
Experiment 2 of salicylic acid treatment of *L. punctata* experiment figure 4c. All wells were treated on the same day and each well is a separate biological replicate. Asterisk marks wells with visible contamination of the buffer with *Pss* B728a.

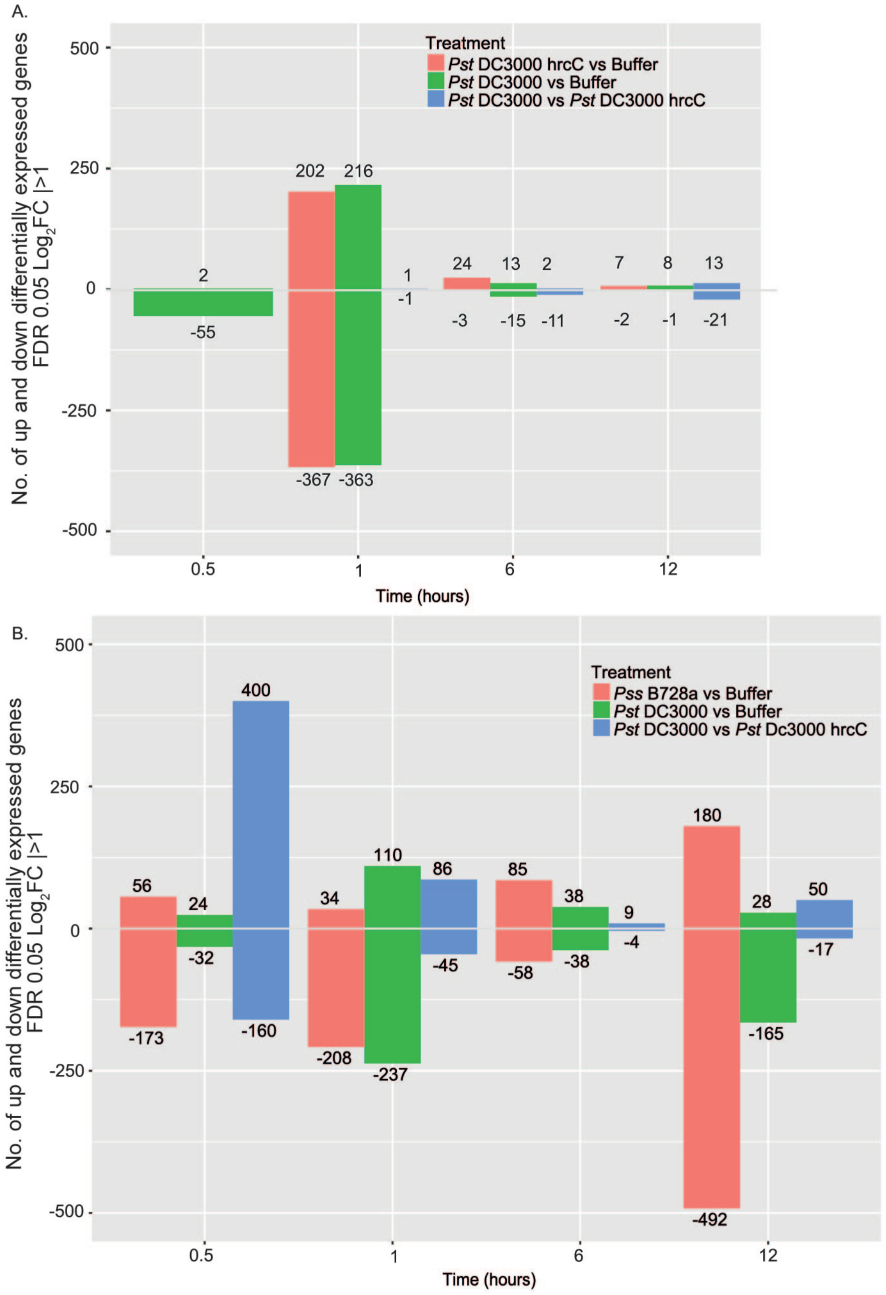


**Sfigure 16. Barchart of number of genes differentially expressed upon bacterial treatments.** Bars indicate no. of up and down differentially expressed genes FDR 0.05 $Log_2FC$ | >1.
A. *S. polyrhiza* differentially expressed genes. B. *L. punctata* differentially expressed genes.

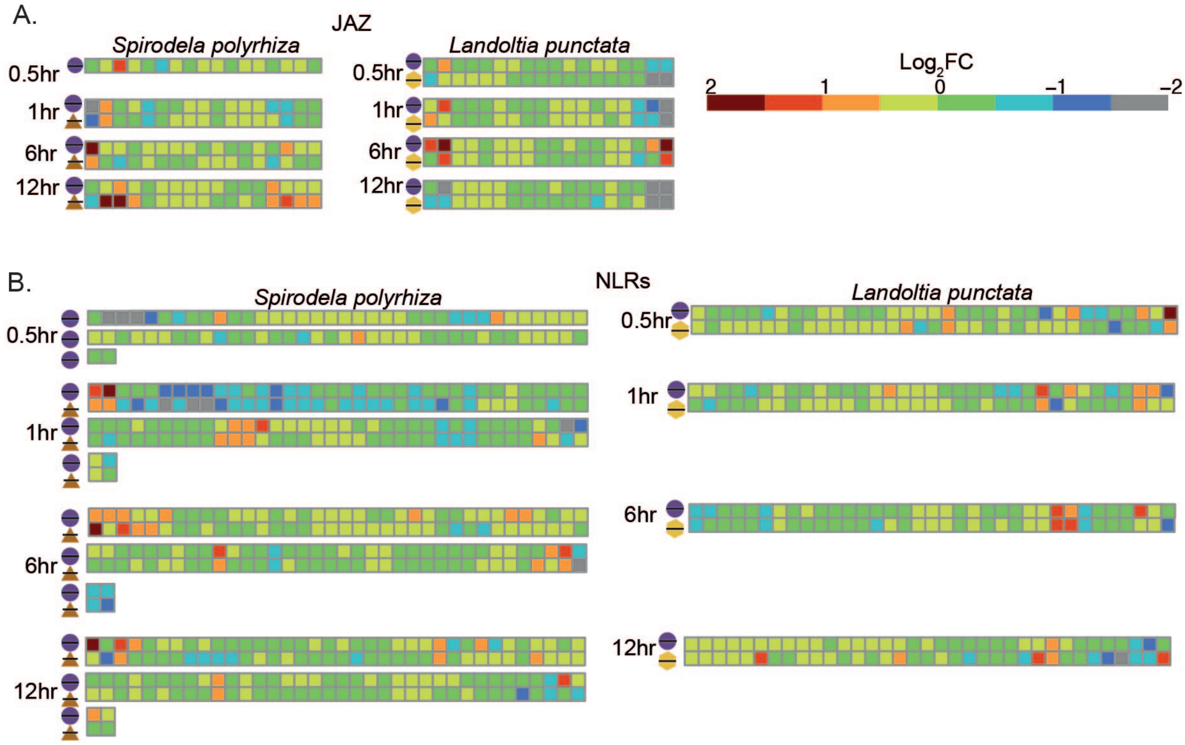


**Sfigure 17. Log$_2$ fold change of selected gene families following bacterial pathogen exposure of duckweeds**. Each square represents a gene with a given domain, the differential expression of the gene is shown by the color of the square. The treatment comparison is indicated by the shapes; black line - buffer, purple circle-*Pst* DC3000, brown triangle - *Pst* DC3000 hrcC and yellow hexagon - *Pss* B728a. JAZ domain containing genes are shown in A and NB-ARC containing genes are indicated in B.

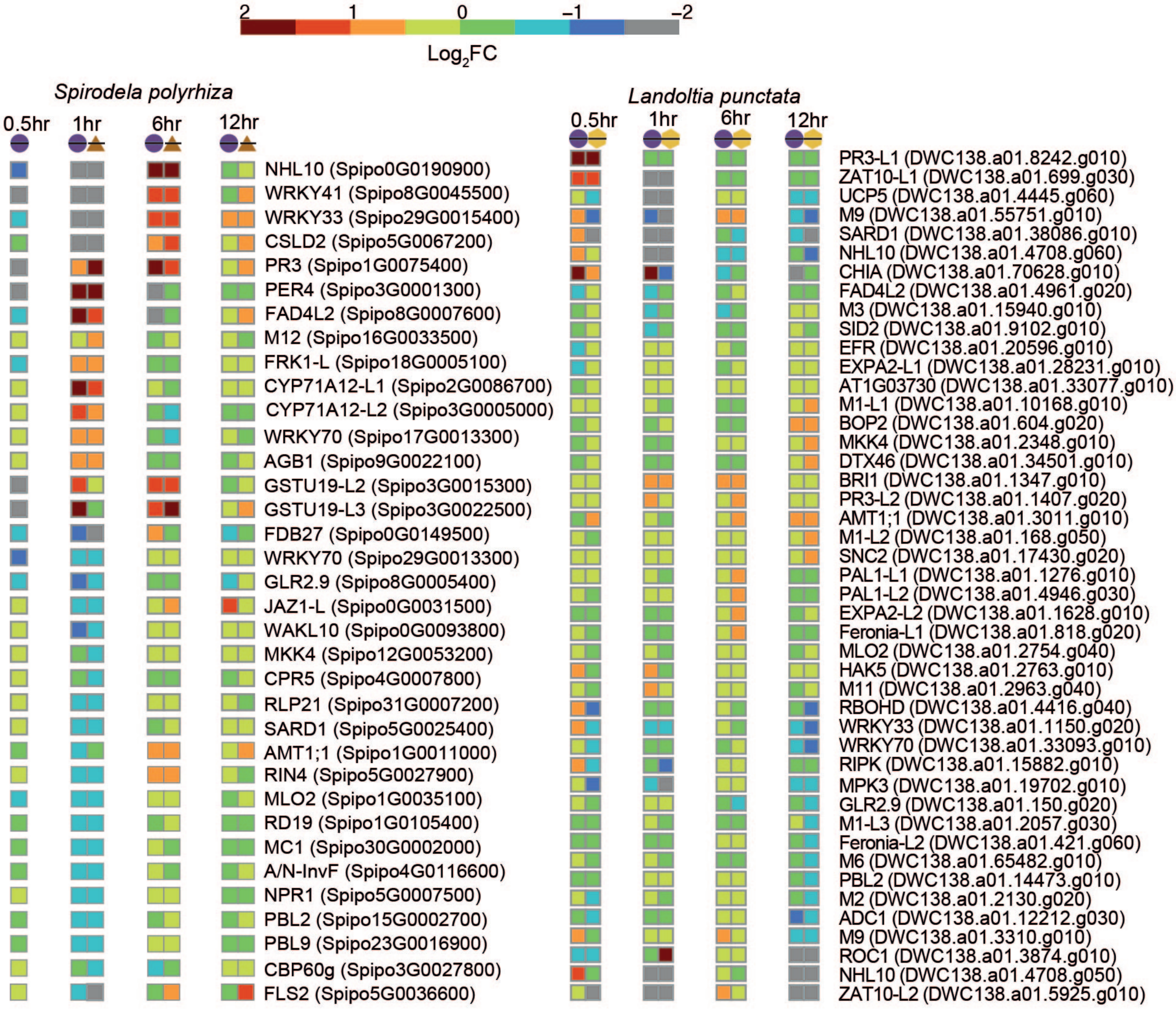


**Sfigure 18. $Log_2$ fold change of homologs of *A. thaliana* bacterial responsive genes.**
Color of square indicates $Log_2$ fold change of homologous genes to *A. thaliana* gene indicated at the end of the row, in brackets is the duckweed gene identified as homologous. The treatment comparison is indicated at the top of the column by the shapes; black line - buffer, purple circle - *Pst* DC3000, brown triangle - *Pst* DC3000 hrcC and yellow hexagon - *Pss* B728a.

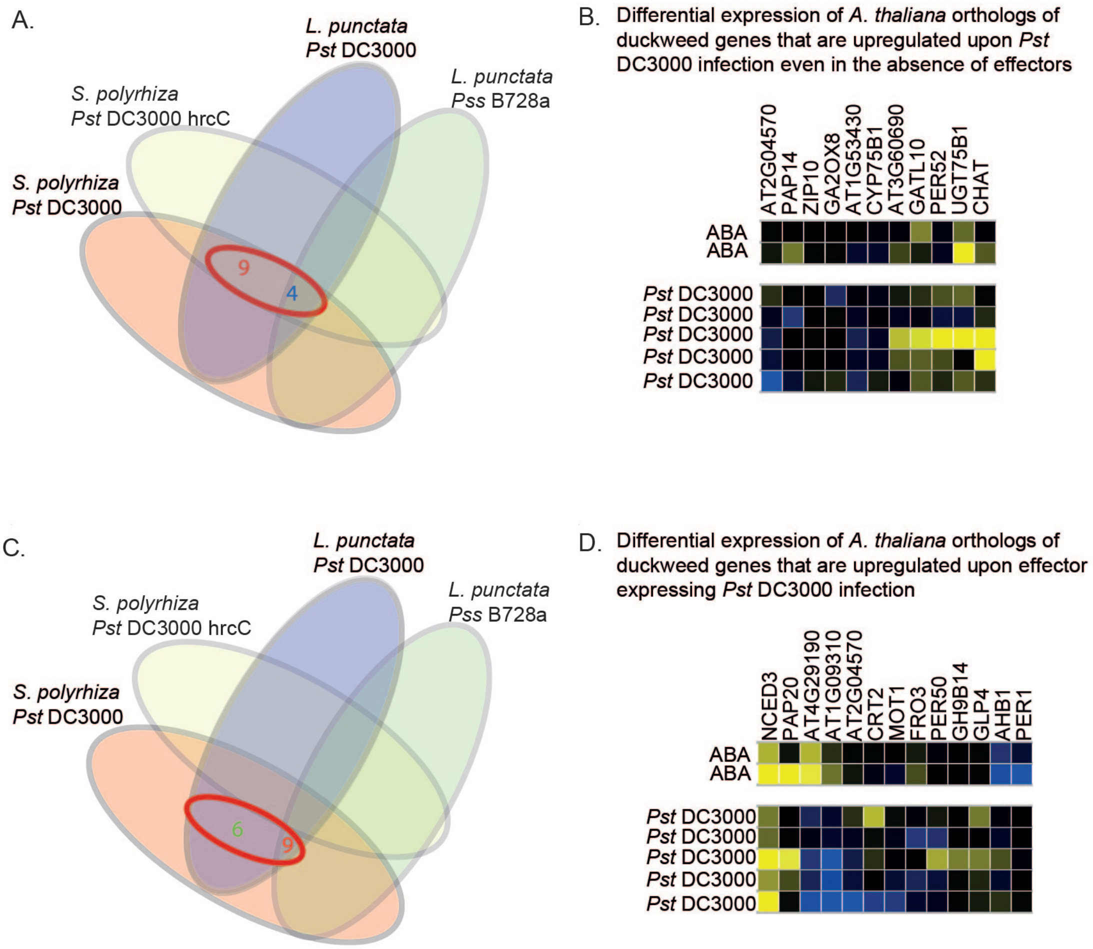


**Sfigure 19. Orthogroups with conserved upregulation upon pathogen treatment of duckweed species and the expression of orthologs in *A. thaliana.***

A. Venn diagram highlighting numbers of orthogroups that are upregulated ($Log_2$ FC >1, FDR < 0.05) in *L. punctata* / *Pst* DC3000 and *S. polyrhiza* / *Pst* D3000 and *Pst* DC3000 hrcC$^{-}$ . B. Microarray differential expression analysis of representatives from *A. thaliana* Col-0 of orthogroups highlighted in A whose upregulation in duckweed is observed in the *Pst* DC3000 hrcC$^{-}$ mutant suggesting their upregulation is independent of effectors. C. Venn diagram highlighting numbers of orthogroups that are upregulated ($Log_2$ FC >1, FDR < 0.05) in *L. punctata* / *Pst* DC3000 and *S. polyrhiza* / *Pst* D3000 but not in *Pst* DC3000 hrcC$^{-}$. D. Microarray expression patterns of subset of genes from C. whose upregulation in duckweed is absent in the *Pst* DC3000 hrcC$^{-}$ mutant suggesting expression is affected by pathogen effectors.

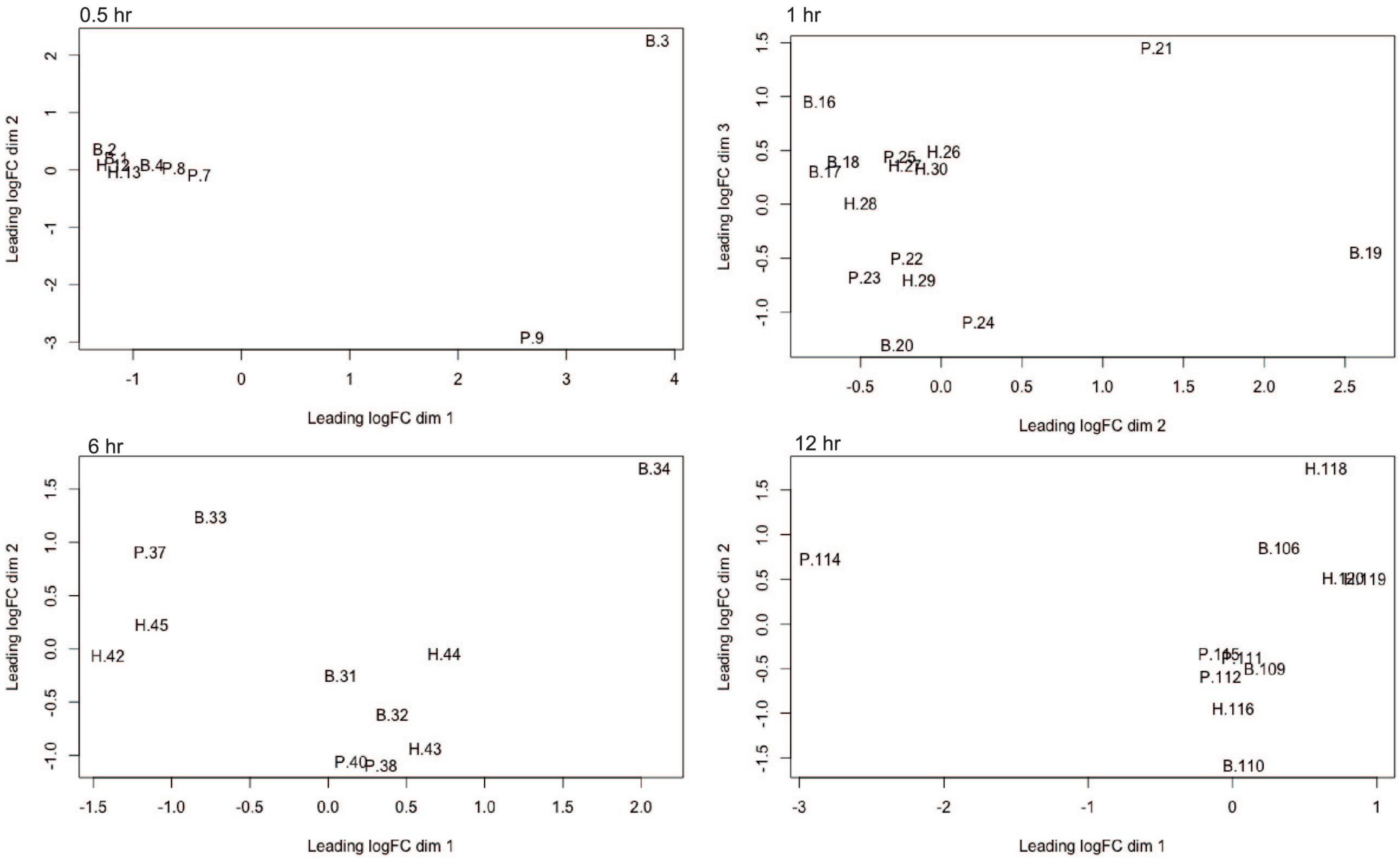


**Sfigure 20. PlotMDS for all *S. polyrhiza* RNAseq samples before outlier removal.**
The PlotMDS dimensions that are shown are those that best separated the treatments. Initial represents treatment condition, B = Buffer, P = *Pst* DC3000 and H= *Pst* DC3000 hrcC.

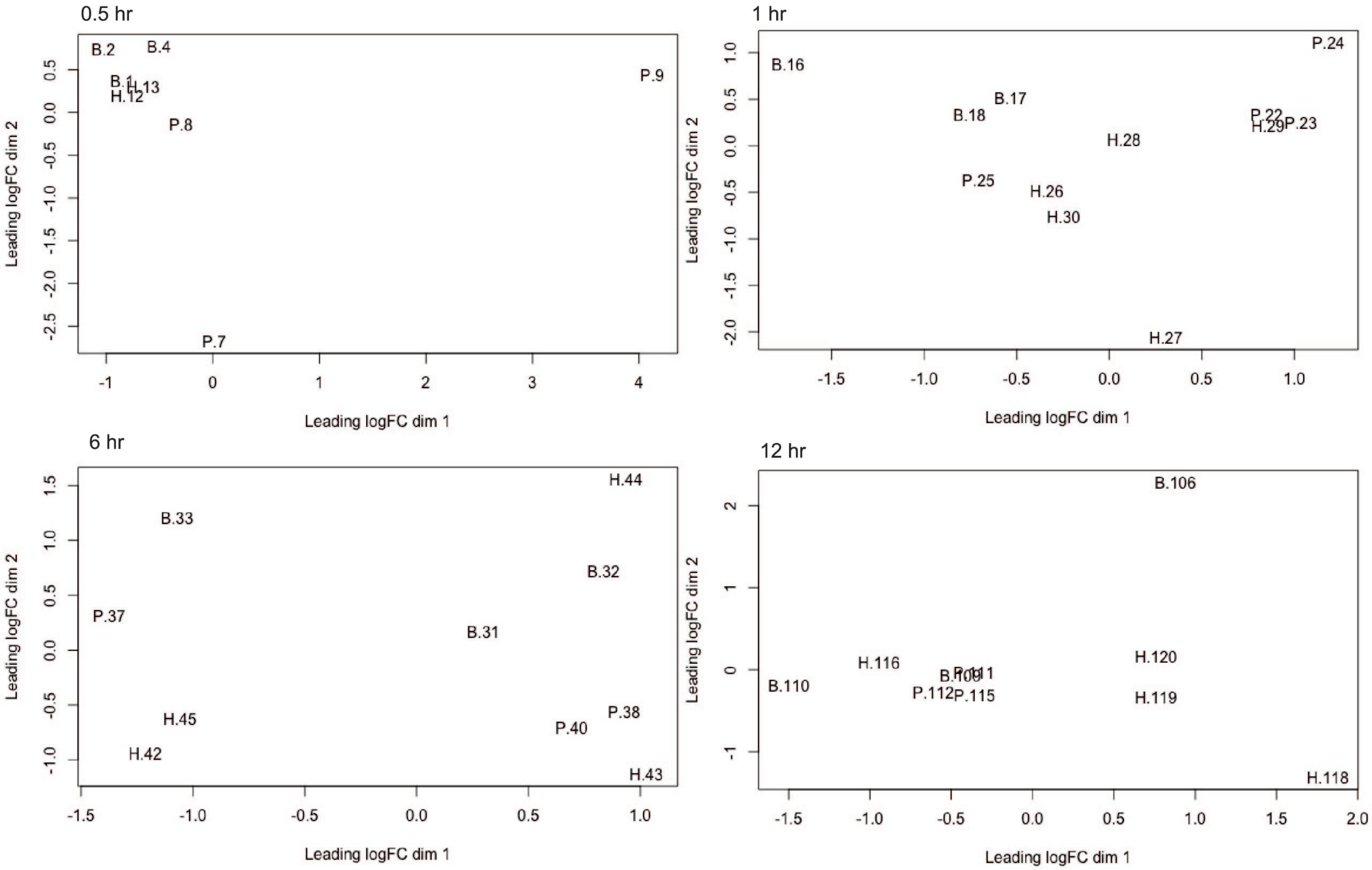


**Sfigure 21. PlotMDS for *S. polyrhiza* RNAseq samples after outlier removal.**
PlotMDS for *S. polyrhiza* RNAseq samples after outlier removal dimensions shown are those that best separates the treatments. Initial represents treatment condition, B = Buffer, P = *Pst* DC3000 and H= *Pst* DC3000 hrcC.

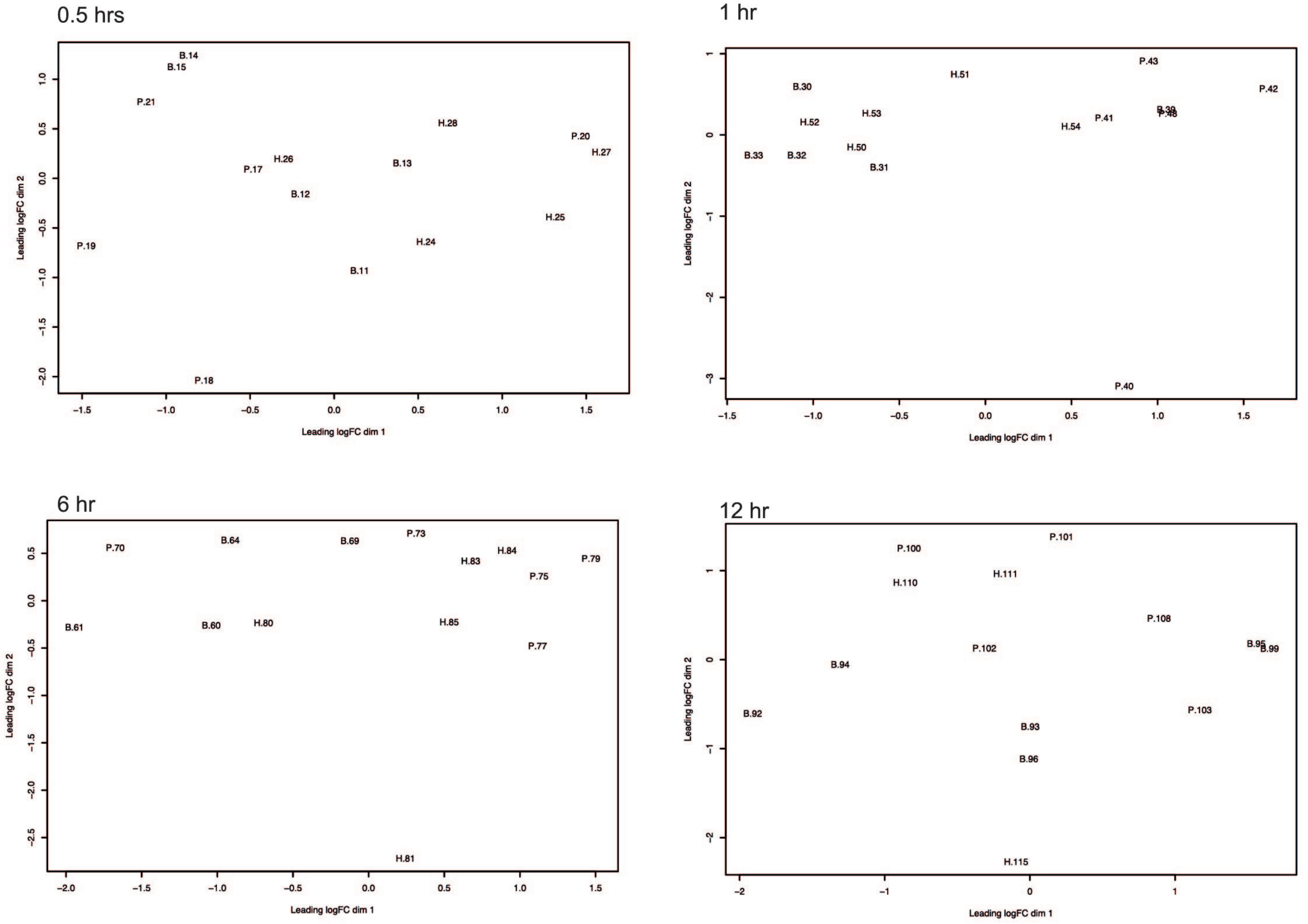


**Sfigure 22. PlotMDS for all *L. punctata* RNAseq samples before outlier removal.**
PlotMDS dim=c(1,2) of all *L. punctata* RNAseq samples for all samples prior to outlier removal. The B prefix indicates buffer H indicates *Pss* B728a and P indicates *Pst* DC3000.

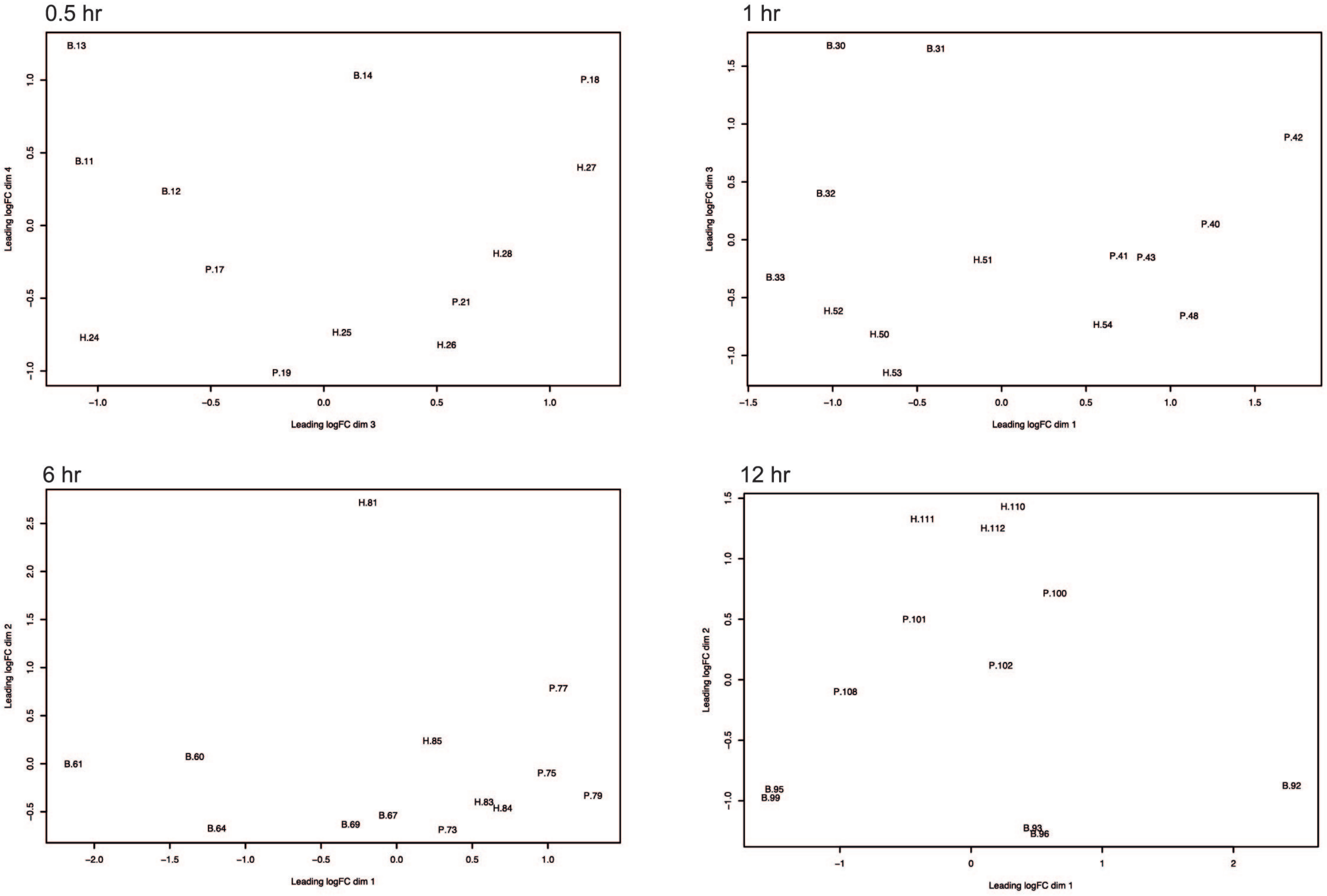


**Sfigure 23. PlotMDS for *L. punctata* RNAseq samples after outlier removal.**
PlotMDS of *L. punctata* RNAseq samples after outlier removal. Dimensions displayed are those that best separate treatments. The B prefix indicates buffer, H indicates *Pss* B728a, and P indicates *Pst* DC3000.
